# Ppard Is Essential in Acceleration of Pancreatic Ductal Adenocarcinoma Development by High-Fat Diet in Mutant *Kras* Mice

**DOI:** 10.1101/2020.12.04.412320

**Authors:** Yi Liu, Yasunori Deguchi, Daoyan Wei, Micheline J. Moussalli, Donghui Li, Huamin Wang, Lovie Ann Valentin, Jennifer K. Colby, Fuyao Liu, Jing Wang, Xiaofeng Zheng, Baoan Ji, James C. Yao, Xiangsheng Zuo, Imad Shureiqi

## Abstract

Pro-obesity high-fat diet is linked with an increased incidence of pancreatic cancers, but the molecular underpinnings of this association remain poorly understood. Here, we report that PPARD is upregulated in pancreatic intraepithelial neoplasia lesions (PanINs) at early stages of pancreatic tumorigenesis in humans and mutant *Kras* mice. Transgenic overexpression of Ppard in pancreatic epithelial cells drastically accelerates the development and progression of pancreatic ductal adenocarcinoma in mutant *Kras* mice when activated by feeding the mice with a high-fat diet or a diet containing GW501516 (50 mg/kg), a selective PPARD agonist. In contrast, pancreatic Ppard genetic deletion significantly suppressed the promotion of pancreatic tumorigenesis by these diets. Mechanistically, we found that this Ppard hyperactivation in pancreatic epithelial cells of mutant *Kras* mice increased production of chemokines and cytokines (e.g., CcI2, CcI4-5, CxcI5 and II6), leading to the robustly increased recruitment of myeloid-derived suppressor cells and macrophages into pancreata, which fostered an immune suppressive microenvironment and subsequently accelerated pancreatic ductal adenocarcinoma development and progression. Our findings demonstrate that PPARD plays an essential role in the promotion of pancreatic tumorigenesis by a high-fat diet. Targeted inhibition of PPARD activation is a potential interventive strategy for pancreatic cancer prevention and therapy.

## INTRODUCTION

Pancreatic ductal adenocarcinoma (PDAC) is a highly lethal disease with a rising incidence ^1^. New and effective preventive and therapeutic strategies are urgently needed. Oncogenic *KRAS* mutation is an initial event in pancreatic carcinogenesis and occurs in more than 92% of pancreatic intraepithelial neoplasia lesions (PanINs) ^2^, which represent early stages of pancreatic tumorigenesis. However, other genetic alterations and inflammatory factors are required to promote *KRAS*-initiated pancreatic tumor progression to PDAC^3^. Thus, identification of critical factors that promote pancreatic tumor progression would lead to the development of new interventions to reduce the incidence and improve dismal prognosis of PDAC.

High-fat diets have been associated with increased risk for human PDAC ^4–6^. In preclinical animal models, high-fat diets promote both chemically-induced and oncogenic *KRAS*–initiated pancreatic carcinogenesis ^7–14^. Fatty acids in high-fat diets act as activating ligands of PPARD, a transcriptional factor that exerts a wide spectrum of effects on important molecular events ^15^. In particular, fatty acid activation of Ppard promotes intestinal tumorigenesis in *Apc*–mutant mice ^16^. However, whether a high-fat diet promotes pancreatic tumorigenesis via upregulation of PPARD remains unknown.

PPARD is upregulated in various major human cancers, including pancreatic cancers ^17–20^. PPARD upregulation in human pancreatic cancers correlates with a higher pathological stage and a higher risk of metastasis ^19^. However, it has been proposed that PPARD upregulation acts differently in different cell types in various organ tumors, promoting tumorigenesis when expressed in stromal cells while inhibiting tumorigenesis when expressed in epithelial cells^19, 21, 22^. In our previous study, we however found that Ppard downregulation in Panc02 mouse pancreatic adenocarcinoma epithelial cells strongly suppressed tumorigenesis ^23^. Others have proposed that PPARD upregulation in pancreatic cancers might be only a stress response indicator rather than being “causally related to the tumor phenotype”, based on *in-vitro* study results showing that a selectively synthetic PPARD agonist, GW501516, inhibited pancreatic cancer cell invasion ^24^. Thus, the mechanistic role of PPARD in pancreatic carcinogenesis remains poorly defined and even controversial.

PPARD synthetic agonists were developed and tested clinically to treat metabolic disorders ^25, 26^. Large pharmaceutical companies however halted the development of PPARD agonists (e.g., GW501516) to treat non-cancerous disorders (e.g., obesity) because of the concerns about their potential carcinogenic effects. Nonetheless, PPARD agonists such as GW501516 are still illicitly sold to athletes to enhance muscle endurance via websites claiming a lack of evidence of harmful effects. The marketing of this agent was further enhanced by major news media coverage of a study published in 2017 showing PPARD’s ability to enhance muscle endurance ^27^. Thus, clarifying the role of PPARD in carcinogenesis is highly critical.

In this study, we aimed to use both genetic and pharmaceutical approaches to determine the essential role of PPARD in pancreatic tumorigenesis, especially when activated by natural fatty acids in high-fat diet–a risk factor for PDAC.

## RESULTS

### PPARD expression is upregulated at early stages of pancreatic tumorigenesis in humans and mice

To determine whether PPARD is upregulated in the earlier stages of pancreatic tumorigenesis, we evaluated PPARD mRNA expression by RNAscope in situ hybridization and evaluated protein expression by immunohistochemical (IHC) staining of PanINs in pancreata from human patients and KC mice (*Pdx1-Cre;* LSL-*Kras^G12D^*), in which the *Kras^G12D^* mutation is targeted to Pdx-1^+^ pancreatic epithelial cells (**sFig. 1A),** along with paired adjacent normal pancreatic tissues. PPARD was weakly expressed in the nuclei of human normal pancreatic tissues, but strongly expressed in the nuclei and cytoplasm of the paired PanINs at both mRNA and protein levels, and this PPARD expression was further increased in PDACs (**Fig. 1A-D** and **sFig. 1B and C**). Similarly, Ppard mRNA expression was upregulated in PanINs compared to the adjacent normal pancreatic tissues in KC mice (**sFig. 1A and Fig. 1E and F**). Histologic examinations of the KC mouse tissues showed that compared to the adjacent normal pancreatic tissues, PanINs had higher expression levels of p-ERK1/2, an indicator for aberrant oncogenic *Kras* activation, as measured by IHC staining, higher levels of acidic mucins, as stained by Alcian blue ^28^; and more pancreatic fibrosis, as shown by Sirius red staining for collagen fibrils ^29^ (**Fig. 1E**).

**Figure 1.**
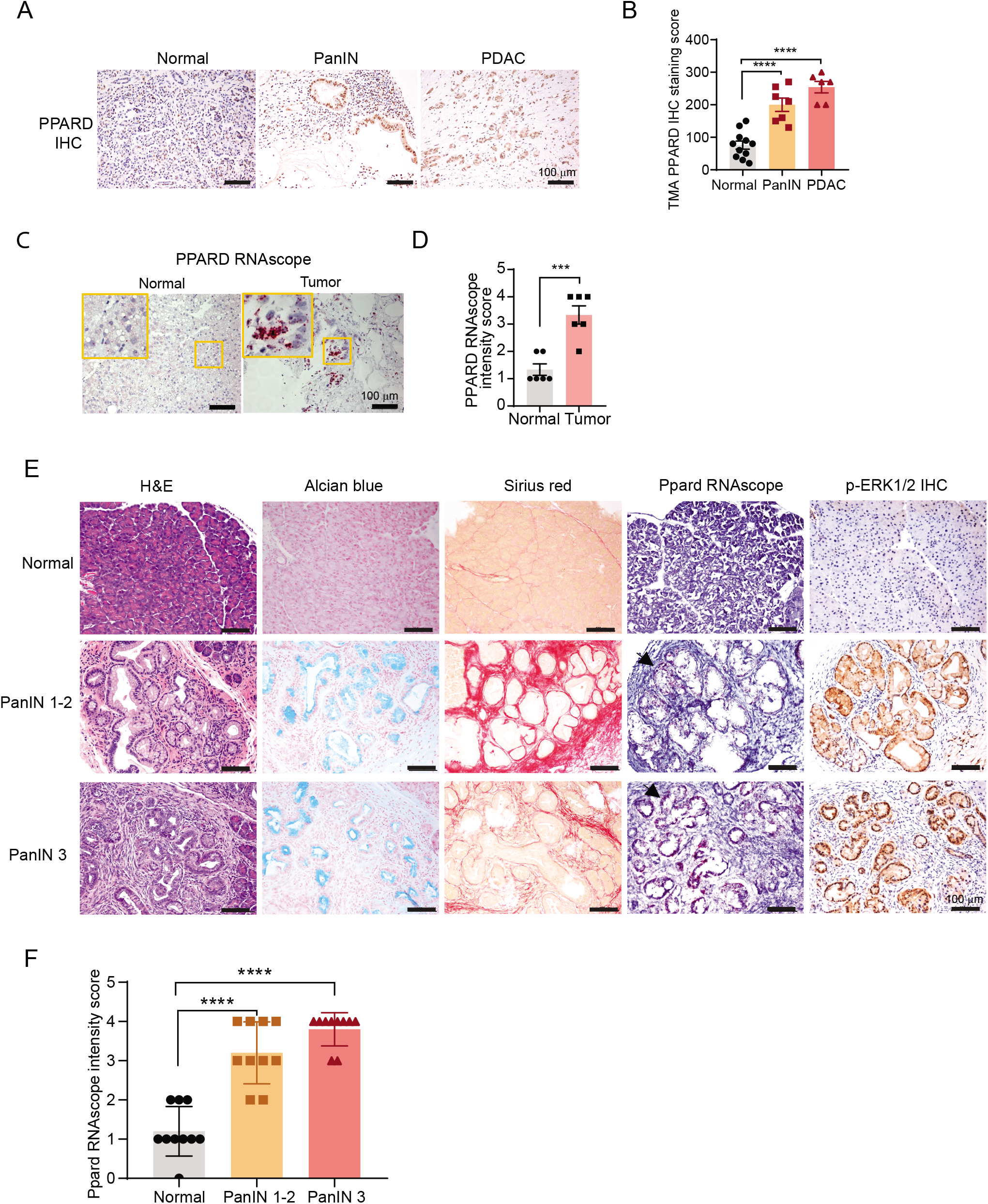
PPARD expression is upregulated in human and mouse pancreatic PanINs. **(A and B)** PPARD protein expression in human normal pancreas, pancreatic intraepithelial neoplasia (PanIN), and pancreatic ductal adenocarcinoma (PDAC) tissues in TMA, measured by immunohistochemistry (IHC). **(A)** Representative images of PPARD IHC staining of the indicated groups of human pancreatic tissue samples. **(B)** PPARD IHC scores of normal (n=11), PanIN (n=7), and PDAC (n=6) pancreata tissues from pancreatic tumor microarray slide (TMA). **(C and D)** PPARD mRNA expression levels in fresh slides of human normal and tumor pancreatic tissues (PanINs) by biopsy, measured by RNAscope in situ hybridization. (**C**) Representative images of PPARD RNAscope in situ hybridization staining of human pancreatic tissue slides. PPARD mRNA stained positive is marked in red. **(D)** The quantitative results of PPARD intensity scores for normal and tumor pancreatic tissues (n=6 fields under 10× bright filed microscope for each group). **(E)** Representative images of hematoxylin and eosin (H&E) staining, Alcian blue staining, Sirius red staining, Ppard RNAscope in situ hybridization, and phospho-ERK1/2 (p-ERK1/2) IHC staining, in normal, PanIN 1-2, and PanIN 3 pancreatic tissues in KC mice. **(F)** The quantitative results of Ppard mRNA intensity scores of normal, PanIN 1-2, and PanIN 3 pancreata tissues in KC mice (n=10 for each group) as described in panel **F**. For **B, D and F,** data are mean ± SEM; \*\*\**P* <.001, \*\*\*\**P* <.0001.

### Transgenic Ppard overexpression in pancreatic epithelial cells augments promotion of oncogenic *Kras^G12D^*–initiated pancreatic tumorigenesis by a high-fat diet

PPARD is upregulated in PanINs of human patients and KC mice (**Fig. 1)**, and this PPARD upregulation increases with the progression from PanIN to PDAC in humans (**Fig. 1A and B**)^19^. Exposure to natural ligands (e.g., fatty acids) and synthetic ligands (e.g., GW501516) of PPARD in the setting of progressive overexpression of PPARD during pancreatic tumor progression would lead to PPARD hyperactivation. We therefore investigated the mechanistic significance of Ppard hyperactivation during carcinogenesis by examining the effects of a high-fat diet or GW501516 in KC mice with transgenic Ppard overexpression in pancreata, to model the progressive Ppard upregulation during pancreatic tumor progression.

To this end, we generated a new mouse model, Panc-Pd (*Pdx1-Cre*; CAG-LSL-*Ppard*), in which Ppard overexpression is targeted to mouse pancreata by a Pdx1 promoter–driven Cre-recombinase system as described in the Methods (**sFig. 2A and B**). Panc-Pd mice had higher pancreatic Ppard expression levels than did their wild type littermates on Western blotting (**sFig. 2C**), suggesting this transgenic mouse model is successful. Transgenic Ppard overexpression alone is not sufficient to induce pancreatic tumorigenesis (data not shown). For pancreatic tumorigenesis study, Panc-Pd mice were then bred with KC mice to generate KC/Pd (*Pdx1-Cre; LSL-Kras^G12D^*; CAG-LSL-*Ppard*) mice for simultaneous expression of Ppard and mutant KRas^G12D^ in the pancreas (**Fig. 2A**). Gross and histologic examinations showed that prior to 12 weeks of age, KC/Pd mice developed very few PanIN lesions, with very low *Kras^G12D^* activity, measured by p-ERK1/2 IHC staining similar to KC mice (data not shown). However, starting from age 12-14 weeks, KC/Pd mice developed larger areas of PanINs than did their KC littermates (**sFig. 2D**), and these lesions expanded more aggressively to larger areas in KC/Pd mice as the mice aged to 20 weeks (**sFig. 2E and F**).

**Figure 2.**
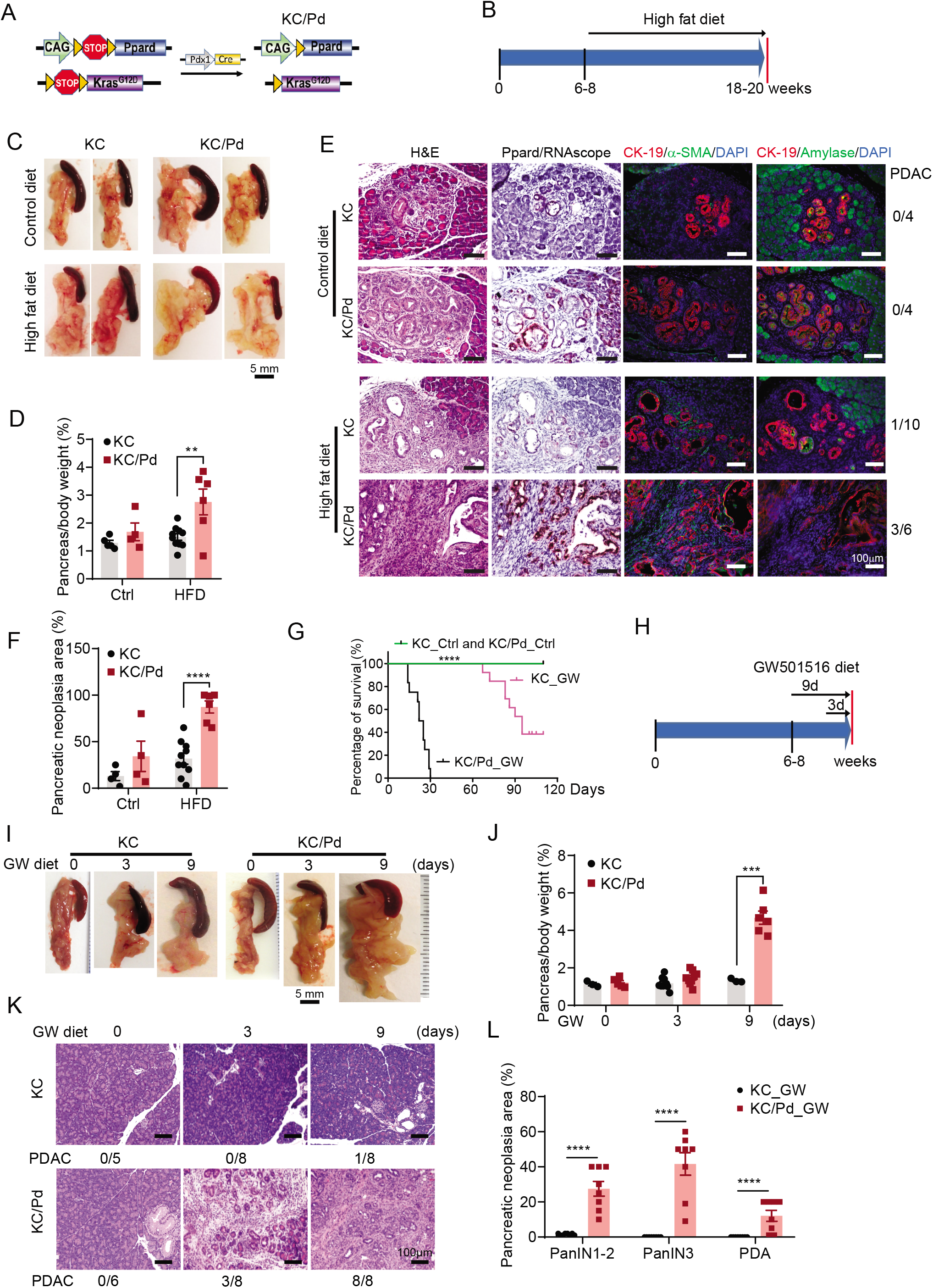
Ppard overexpression in pancreatic epithelial cells augments promotion of pancreatic tumorigenesis in KC mice by a high-fat or a GW diet. **(A)** Schematic diagram for the generation of KC/Pd mice. **(B-F)** KC and KC/Pd mice at 6-8 weeks old fed either a high-fat diet (60% kcal from fat) or a control diet (10% kcal from fat) for 12 weeks were euthanized, and pancreata were photographed, weighed, and harvested for gross and histologic characterization (n=4-10 per group). **(B)** Schematic diagram for timeline of a high-fat diet treatment. **(C)** Representative images of pancreata from KC and KC/Pd mice fed the high-fat or the control diet. **(D)** The weight ratio of pancreas to body per mouse for the indicated mouse groups. **(E)** Representative images of H&E staining, Ppard RNAscope in situ hybridization, CK-19 and α-SMA co-immunofluorescence staining, and CK19 and amylase co-immunofluorescence staining of the pancreata from the indicated mouse groups. **(F)** Percentage of pancreatic neoplastic area per mouse. **(G)** Survival of KC or KC/Pd mice on a GW501516 diet. KC or KC/Pd mice at age 6-8 weeks were fed a diet containing GW501516 (GW, 50 mg/kg) or the same diet except without GW501516 as a control (Ctrl) (n=12-15 mice per group). The survival time was calculated by the Kaplan-Meier method and compared between groups using the log-rank test. \*\*\*\**P* <.0001 **(H-L)** KC or KC/Pd mice at 6-8 weeks were fed a diet containing GW (50 mg/kg) or a control diet (Ctrl) for 0, 3, or 9 days, then euthanized and analyzed for pancreatic tumorigenesis (n=5-8). **(H)** Schematic diagram for timeline of GW diet treatment. **(I)** Representative images of pancreata from KC and KC/Pd mice fed the GW diet for the indicated days. **(J)** The weight ratio of pancreas to body per mouse for the indicated mouse groups**. (K)** Representative images of H&E-stained sections of pancreata from the indicated mice. **(L)** Percentage of pancreatic neoplastic area per mouse for PanIN 1-2, PanIN 3, and PDAC in KC or KC/Pd mice fed the GW diet for 9 days. For **D, F, J**, and **L**, Data are mean ± SEM; \*\**P* <.01, \*\*\**P* <.001, \*\*\*\**P* <.0001.

Next, we investigated whether a high-fat diet, a natural activator of Ppard, promotes pancreatic tumorigenesis through activation of Ppard by feeding KC and KC/Pd mice at age 6-8 weeks with either a high-fat (60% kcal from fat, Envigo) or a control diet (10% kcal from fat, Envigo) for 12 weeks (**Fig. 2B**). In KC mice, high–fat diet increased sizes and weights of both pancreata and abdominal fats, but ratios of pancreas to body weights remained unchanged (**Fig. 2C and D**, **sFig. 2G**). In contrast, the high-fat diet failed to increase abdominal fat and body weights in KC/Pd mice (**sFig. 2G)**. Nevertheless, KC/Pd mice on the high fat diet had marked increases of acinar cell dedifferentiation to duct-like cells and fibrosis development as measured by immunofluorescence staining (IF) of amylase, cytokeratin-19 (CK-19) and α-smooth muscle actin (α-SMA) (**Fig. 2E**), and pancreatic neoplastic lesions, compared to KC mice on the high fat diet and KC/Pd and KC mice on the control diet (**Fig. 2C-F**). These high-fat diet–induced tumorigenic effects were markedly augmented in KC/Pd mice (**Fig. 2C-F**). Interestingly, a high-fat diet also increased Ppard expression levels in pancreata of both KC and KC/Pd mice (**sFig. 2H**). These data suggest that PPARD hyperactivation mediated protumorigenic effects of a high-fat diet during pancreatic tumorigenesis.

To further investigate the mechanistic role of Ppard hyperactivation in the promotion of pancreatic tumorigenesis by a high-fat diet, we used PPARD-specific synthetic ligand GW501516 in the same models of pancreatic tumorigenesis. KC and KC/Pd littermates at 6-8 weeks old were randomly assigned either a diet containing GW501516 (GW, 50 mg/kg, Envigo) or the same diet except without GW (control). Unexpectedly, KC/Pd mice died after 15-30 days of GW diet treatment, and more than 50% of KC mice died after 90 days of GW diet treatment, but none of sex- and age-matched KC or KC/Pd mice fed the control diet died within the observed time period (115 days) (**Fig. 2G**). We next investigated the cause of those dramatical survival differences in KC and KC/Pd mice at around 6-8 weeks of age by feeding them the GW diet for 0 days (i.e., control diet for 9 days), 3 days (control diet for 6 days and then GW diet for 3 days), and or 9 days (GW diet for all 9 days) and then performing gross and histologic examination of their pancreata (**Fig. 2H**). While KC mice showed minimal PanINs after 9 days of GW diet, 100% of KC/Pd mice developed PanINs with severe pancreatitis and significant acinar cell loss and 37.5% (3/8) of KC/Pd developed PDA after 3 days of the GW diet treatment, and 100% (8/8) of KC/Pd mice exhibited diffuse pancreatic tumors, including advanced PanIN 3 and PDAC development, after 9 days of the GW diet (**Fig. 2I-L**). Furthermore, we found that the KC/Pd mice that died after receiving GW diet for 15-30 days developed advanced pancreatic tumors with almost 100% acinar cell loss, severe pancreatitis, and fibrosis (**sFig. 2I**). However, Panc-Pd mice fed the GW diet for 100 days failed to develop pancreatic neoplastic lesions (data not shown). These data demonstrated that Ppard hyperactivation dramatically promoted high-fat diet–induced pancreatic tumorigenesis in mutant *Kras* mice.

### Ppard genetic deletion in pancreatic epithelial cells inhibits promotion of oncogenic *Kras^G12D^*–initiated pancreatic tumorigenesis by a high-fat diet

To assess whether Ppard hyperactivation in pancreatic epithelial cells is essential to the promotion of oncogenic *Kras^G12D^*–initiated pancreatic tumorigenesis by a high-fat diet, we generated a new mouse model with specific Ppard genetic deletion and *Kras^G12D^* mutation in Pdx1^+^ pancreatic epithelial cells by breeding KC mice with Ppard conditional knockout (KO) (Ppard-flox) mice (KC/PdKO, **Fig. 3A**). As expected, KC/PdKO mice have extremely low mRNA levels of Ppard in their pancreata **(Fig. 3B**). KC and KC/PdKO mice at 6-8 weeks were then fed either a high-fat or a control diet for 26 weeks (**Fig. 3C**). We found that in KC mice, the high-fat diet significantly increased sizes and weights of pancreata (**Fig. 3D and E**) and pancreatic neoplastic lesions (**Fig. 3F and G**), compared to the control diet, whereas all these pro-tumorigenic effects were significantly decreased in the KC/PdKO littermates on the high-fat diet compared to KC mice on the high-fat diet (**Fig. 3D-G**). Furthermore, PDA was observed in 14.3% (1/7) and 44.4% of KC mice fed a control and a high-fat diet respectively, but not in any of KC/PdKO mice fed either a control or a high-fat diet. Our results indicated that Ppard activation is required for the the promotion of pancreatic tumorigenesis by a high-fat diet.

**Figure 3.**
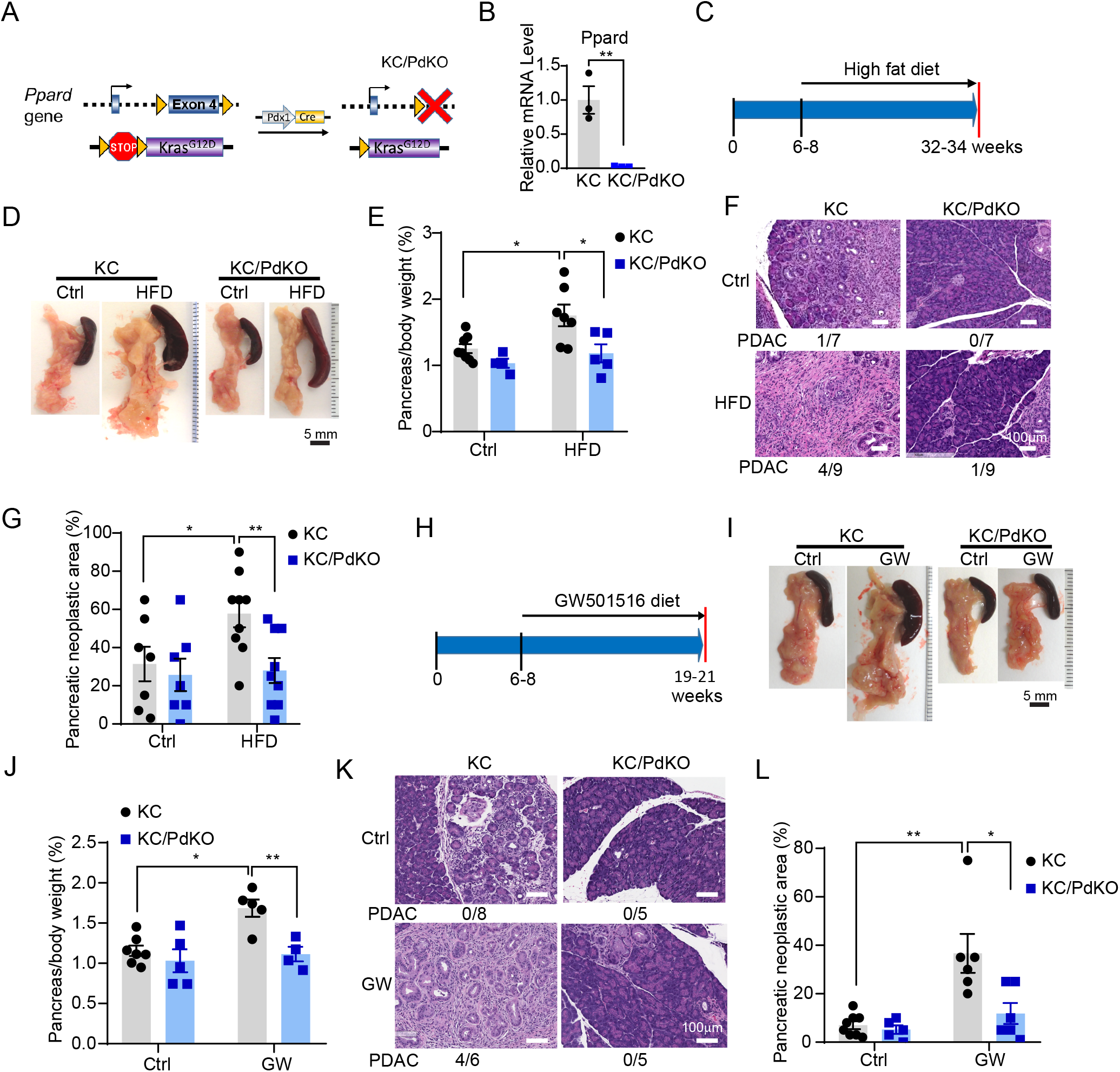
Ppard genetic deletion in pancreatic epithelial cells suppresses pancreatic tumorigenesis in KC mice on a high-fat or a GW diet. **(A)** Schematic diagram of the generation of KC/PdKO mice. **(B)** Pancreatic Ppard mRNA expression in pancreatic tissues from the indicated mice, measured by qRT-PCR (n= 3 per group). **(C-G)** KC and KC/PdKO mice at 6-8 weeks (n=5-8 per group) were fed either a high-fat (HFD) or a control diet (Ctrl) for 26 weeks and then euthanized. Pancreata were photographed, weighted, harvested, and further analyzed. **(C)** Timeline for the mice with a high-fat diet treatment. **(D)** Representative photos of pancreata for the indicated mouse groups. **(E)** The weight ratio of pancreas to body per mouse for the indicated mouse groups. **(F)** Representative pancreas images of H&E staining for the indicated mouse groups. **(G)** The percentage of pancreatic neoplastic area per mouse for the indicated mouse groups. **(H-L)** KC and KC/PdKO mice at 6-8 weeks were fed either a GW (50 mg/kg) or a Ctrl diet for 13 weeks and then euthanized (n=5-8 mice per group). Pancreata were photographed, weighted, harvested, and further analyzed. **(H)** Timeline for the mice with the GW diet treatment. **(I)** Representative photos of pancreata for the indicated mouse groups. **(J)** The weight ratio of pancreas to body per mouse for the indicated mouse groups. **(K)** Representative pancreas images of H&E staining for the indicated mouse groups. **(L)** Percentage of pancreatic neoplastic area per mouse for the indicated mouse groups. For **B, E, G, J**, and **L**, Data are mean ± SEM; \**P* <.05, \*\**P* <.01.

To further characterize the function of Ppard in Kras-initiated pancreatic tumorigenesis, we investigated the effect of Ppard-KO in pancreatic epithelial cells on the ability of the specific Ppard agonist GW501516 to promote pancreatic tumorigenesis in KC/PdKO mice. Treatment with a GW diet also failed to significantly increase pancreatic weights or neoplastic lesions in KC/PdKO mice, even with 13 weeks of treatment (**Fig. 3H**), but drastically and significantly promoted pancreatic tumorigenesis in sex- and age-matched KC mice (**Fig. 3I-L and sFig. 2J**), and 66.7% (4/6) of the GW diet-KC mice developed PDA, demonstrating that the effect of GW501516 on promoting pancreatic tumorigenesis is dependent on and specific to Ppard.

### Ppard hyperactivation recruits immuno-suppressive inflammatory cells that promote pancreatic tumorigenesis

We next examined the mechanisms by which Ppard hyperactivation dramatically promoted pancreatic tumorigenesis by performing comparative RNA–sequencing (RNA-Seq) transcriptome studies of pancreata from KC and KC/Pd mice at 6-8 weeks fed a GW diet (50mg/kg) for 3 days. GW diet–treated KC/Pd and KC mice had distinctive differential expression patterns (**Fig. 4A**). Gene Set Enrichment Analyses of the differentially expressed genes (cut off criteria: pAdj<.05) using Hallmark gene sets showed that the top enriched pathways including epithelial-mesenchymal transition, IL6-JAK-STAT3, Inflammatory response, and KRAS signaling (**Fig. 4B and C, sFig. 3A),** an alternative GSEA analysis using Gene Ontology (GO) gene sets further revealed strong connections between these differentially expressed genes and immune system regulations **(sFig. 3B**).

**Figure 4.**
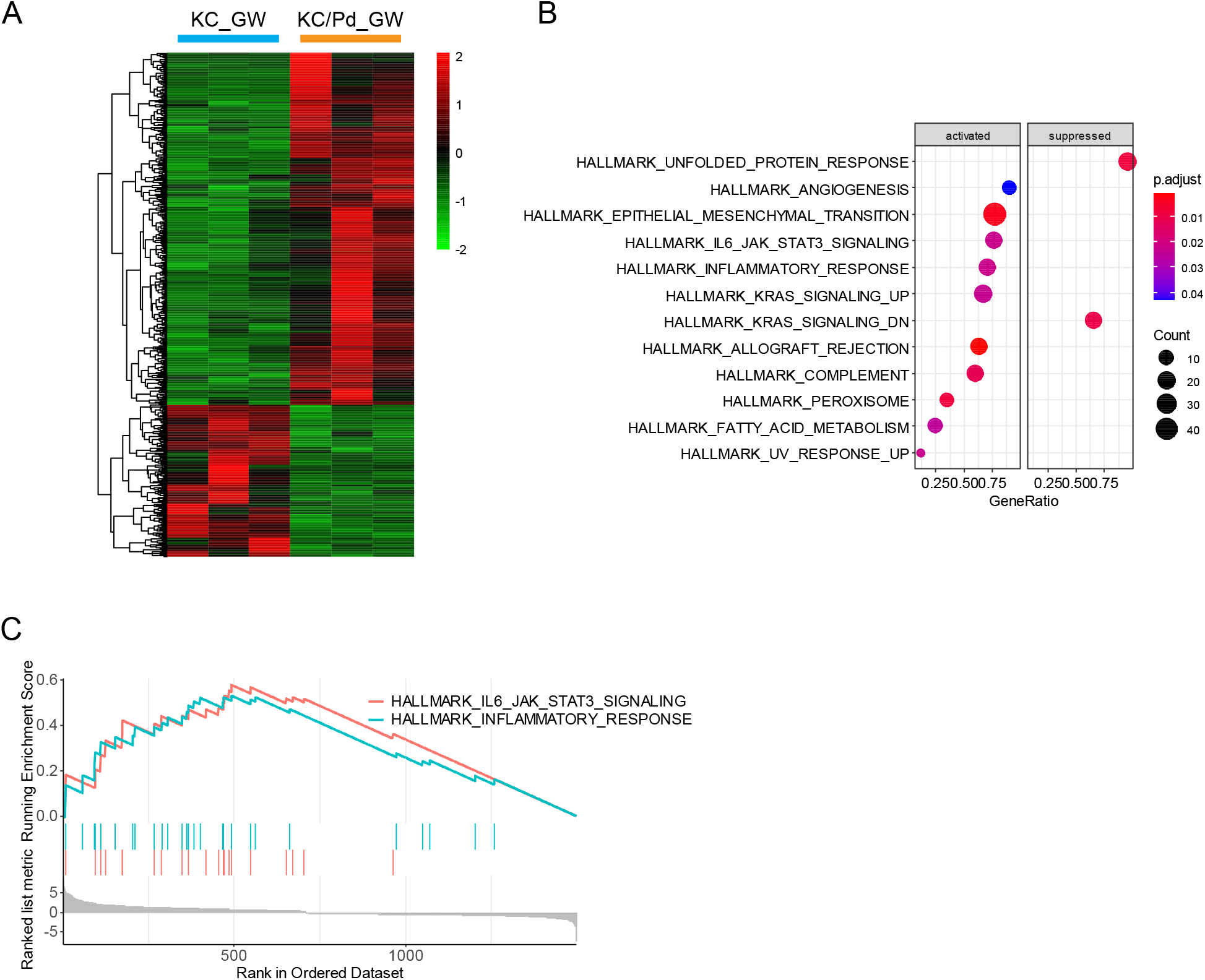
Ppard overexpression in pancreatic epithelial cells promotes inflammatory signaling pathways in KC mice on a GW diet. **(A-C)** KC and KC/Pd mice at 6-8 weeks of age were fed a GW diet (50 mg/kg) for 3 days and then euthanized. The total RNA of pancreata tissues from these mice was extracted for RNA–sequencing transcriptome profile analyses (n=3-4 mice per group). **(A)** Heatmap of differentially expressed genes (DEGs) between a GW diet–treated KC (KC_GW) and a GW diet–treated KC/Pd (KC/Pd_GW) mouse groups. **(B)** Pathway enrichment results for KC_GW and KC/Pd_GW mice obtained by gene set enrichment analyses in R package ClusterProfiler, pAdjCutoff=0.05, gene sets=MSigDB category “Hallmark gene sets”. **(C)** The map of gene enrichment plots of two top inflammatory pathways from the top five signaling pathways shown in panel **B**.

The potential tumorigenic role of these Ppard-mediated pathways, along with the known importance of inflammation in the promotion of oncogenic *Kras*-initiated pancreatic tumorigenesis by a high-fat diet ^12^, led us to investigate the role of Ppard in regulating pancreatic immune and inflammatory responses. We therefore profiled immune and inflammatory cells in the pancreata of KC and KC/Pd mice fed a GW diet (50 mg/kg) for 0, 3, and 9 days as described above. The GW diet markedly increased accumulation of pancreatic myeloid cells (CD11b^+^), including myeloid-derived suppressor cells (MDSCs) and macrophages (CD45^+^CD11b^+^F4/80^+^), in KC/Pd mice compared to their sex- and age-matched KC littermates at 9 days of GW treatment (**Fig. 5A-F, sFig. 4A**). Specifically, among MDSCs, the GW diet increased polymorphonuclear MDSCs (PMN-MDSCs, CD45^+^CD11b^+^Ly6C^+^Ly6G^+^) at 3 days of treatment and continued the increase through 9 days of treatment, while the increase in monocytic MDSCs (M-MDSCs, CD45^+^CD11b^+^Ly6C^hi^ Ly6G^-^) plateaued after 3 days of treatment (**Fig. 5D and E, sFig. 4A**). Interestingly, KC/Pd mice fed the GW diet for 3 days had higher levels of macrophages with Ly6C^low^ cells that had low expression of surface marker MHC-II, an indicator of their polarization into M2-like tumor-associated macrophages (M2-TAM, CD45^+^CD11b^+^F4/80^+^Ly6C^low^MHC-II^low^)^30, 31^, compared to their KC littermates fed the same GW diet; and these effects were augmented after GW treatment for 9 days (**Fig. 5G**). In contrast, the GW diet markedly decreased levels of T cells (CD3^+^CD8^+^ and CD3^+^CD4^+^) in KC/Pd mice, compared to the KC littermates at 9 days of GW treatment (**Fig. 5A and H, sFig. 4B)** but had no significant effects on B cell accumulation (**sFig. 4C)**.

**Figure 5.**
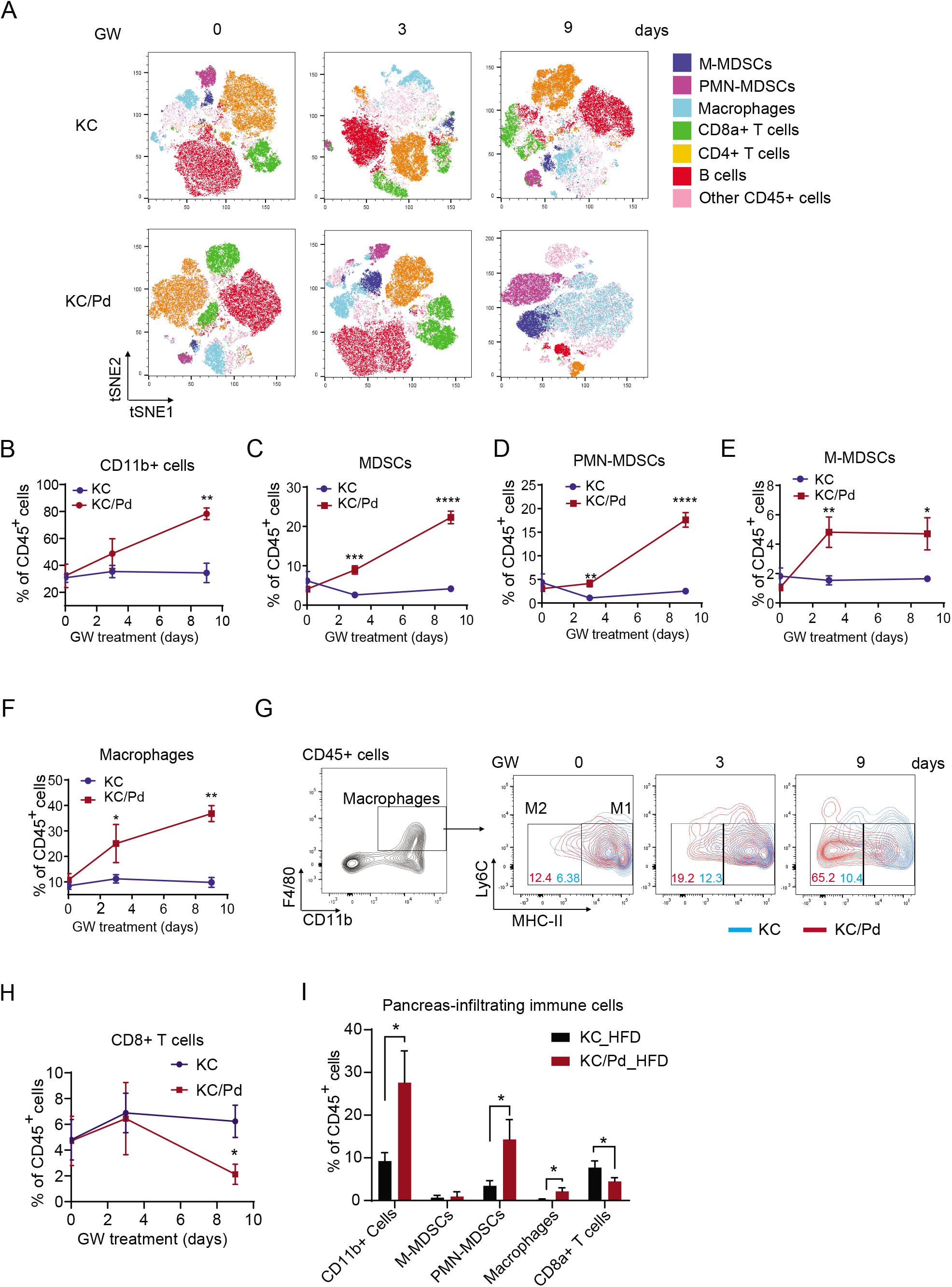
Ppard hyperactivation recruits immunosuppressive inflammatory cells into local pancreata to promote pancreatic tumorigenesis in KC mice. **(A-H)** KC and KC/Pd mice at 6-8 weeks were fed a GW diet (50 mg/kg) for 0, 3, or 9 days and then euthanized (n=4-6 per group). Pancreatic tissues were examined for quantification of infiltrating immune cells measured by flow cytometry. **(A)** Representative immune cell profiling by FlowJo-tSNE (t-distributed stochastic neighbor embedding) of pancreas-infiltrating CD45^+^ cells, including CD11b cells (CD11b^+^), macrophages (CD11b^+^F4/80^+^), total myeloid-derived suppressor cells (MDSCs), monocytic (M) MDSCs (CD11b^+^Ly6C^hi^Ly6G^-^), polymorphonuclear (PMN) MDSCs (CD11b^+^Ly6C^+^Ly6G^+^), and CD3^+^CD4^+^ and CD3^+^CD8^+^ T cells. **(B-F)** Quantitative results for individual cell types including CD11b^+^ **(B),** MDSCs **(C)**, PMN-MDSCs **(D)**, M-MDSCs **(E)**, and macrophages **(F)** for the indicated mouse groups. **(G)** Representative flow cytometry images and quantitative results of the pancreas-infiltrating Ly6C^low^ inflammatory macrophages with low MHC-II expression (M2) for the KC and KC/Pd mice fed the GW diet for 0, 3, or 9 days. **(H)** Quantitative results of CD8^+^ T cells in pancreas-infiltrating CD45^+^ cells for the indicated mouse groups. Data are mean ± SEM; \**P* <.05, \*\**P* <.01, \*\*\**P* <.001, \*\*\*\**P* <.0001, compared to KC mice with the same diet treatment. **(I)** Quantitative results of indicated immune cells in pancreas-infiltrating CD45^+^ cells in KC and KC/Pd mice fed a high fat diet for 12 weeks, as described in **Fig. 2B-F**. Data are mean ± SEM; \**P* < 0.05.

Then, to determine the effects of Ppard hyperactivation via its natural ligands on immune and inflammatory cell recruitment, we performed the same profiling of pancreas-infiltrating immune cells in KC and KC/Pd mice fed a high-fat diet for 12 weeks. We also found that pancreata of KC/Pd mice had significantly higher accumulation of CD45^+^CD11b^+^ cells, PMN-MDSCs, and macrophages, but lower accumulation of CD3^+^CD8^+^ T cells, than the pancreata of KC mice did (**Fig. 5I**).

Our results demonstrated that Ppard hyperactivation via GW treatment or a high-fat diet in KC/Pd mice, increased the recruitment of MDSCs and macrophages into the pancreas, thus orchestrating immuno-suppressive local microenvironment that promoted pancreatic tumorigenesis.

### Ppard hyperactivation increases production of multiple chemokines and cytokines in pancreata in KC mice

To investigate the mechanisms by which Ppard hyperactivation increased the recruitment of inflammatory immunosuppressive cells (MDSCs and macrophages) into the pancreata, we performed screening for a panel of chemokines and cytokines in pancreata of KC and KC/Pd fed a GW diet for 0, 3, or 9 days or fed a high-fat or control diet for 12 weeks as described above. For this screening, we used the LEGEND plex Mouse Proinflammatory Chemokine Panel, which simultaneously quantifies 13 major inflammatory chemokines and the LEGEND plex Mouse Proinflammatory Cytokine Panel, which simultaneously quantifies 13 major inflammatory cytokines, as described before ^32^. KC/Pd mice fed the GW diet for 3 days had significantly higher production of 11 of the 13 screened chemokines (Ccl2-5, Ccl11,Ccl17,Ccl20, Ccl22,Cxcl1, Cxcl9, and Cxcl13) than did KC mice on the GW diet for 3 days, and these 11 upregulated chemokines and Cxcl5 in pancreata of KC/Pd mice further increased at 9 days on the GW diet, compared to their KC littermates on the GW diet for 9 days or compared to the KC/Pd mice on the GW diet for 3 days (**Figs. 6A-D, sFig. 5A-D**). Similarly, KC/Pd mice fed high-fat diet for 12 weeks also had significantly higher production of 5 of 13 screened chemokines (Ccl2, Ccl4-5, Ccl17, and Cxcl5) than did the KC littermates on the high-fat diet for 12 weeks or than did the KC/Pd mice on the control diet (**Fig. 6E-H, sFig. 5E and F**).

**Figure 6.**
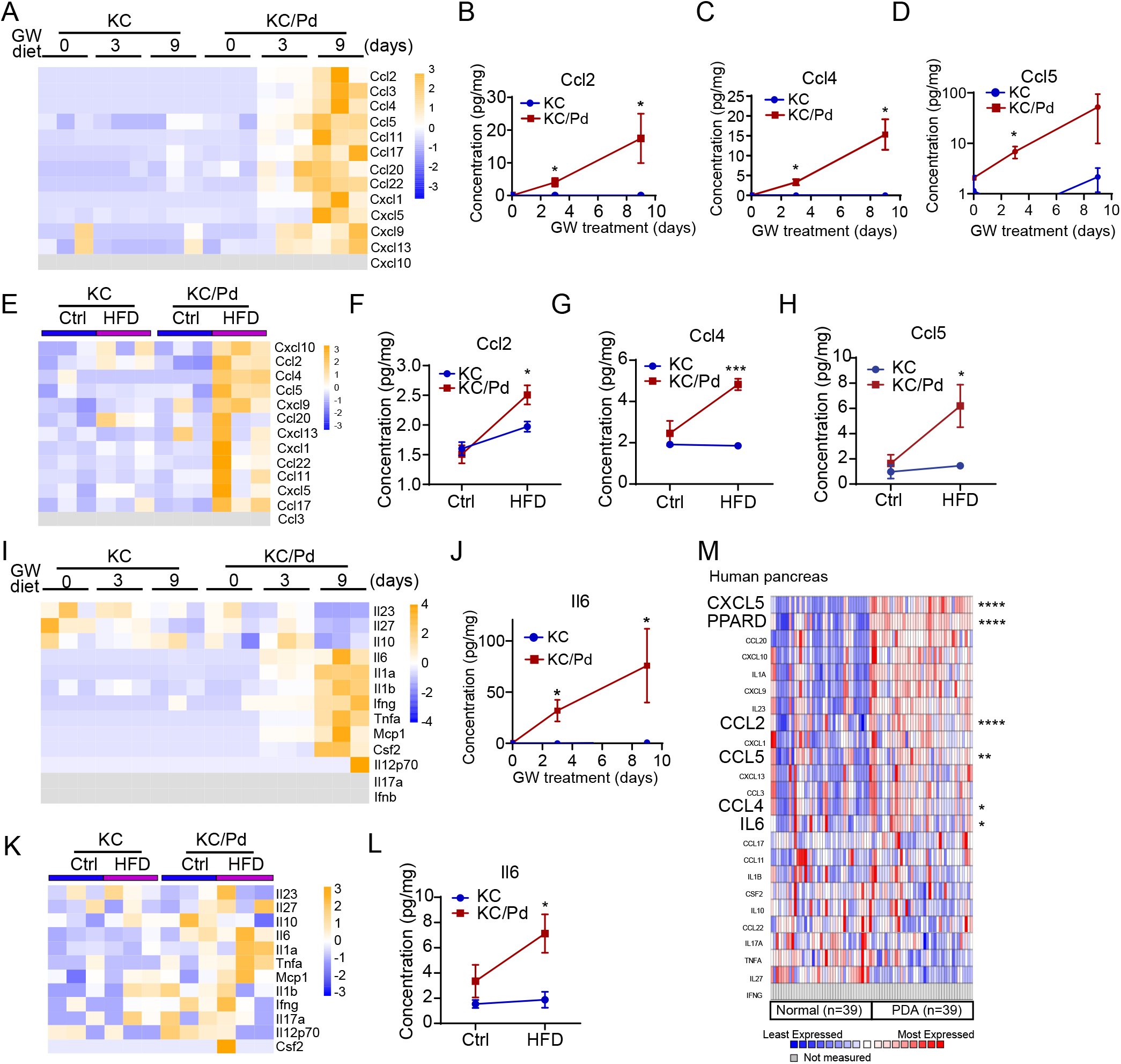
Ppard hyperactivation increases production of multiple pro-inflammatory chemokines and cytokines in pancreata of KC mice. **(A-D)** KC and KC/Pd mice at 6-8 weeks were fed GW (50 mg/kg) for 0, 3, or 9 days and then euthanized. Pancreata were harvested and processed for measurements of a panel of 13 chemokines by using BioLegend’s LEGENDplex bead-based immunoassays (n=3-5 per group). **(A)** Heatmaps of 13 screened chemokines from 3 representative mice per group are shown. Shown are quantitative results for chemokines related to MDSCs’ mobilization and recruitment, including Ccl2 **(B)**, Ccl4 **(C)**, and Ccl5 **(D)**. **(E-H)** KC and KC/Pd mice at 6-8 weeks fed either a HFD or a Ctrl diet for 12 weeks were euthanized. Pancreata were harvested and processed for measurement of the same panel of chemokines as described in **Fig. 6A-D** (n=3-4 per group). (**E**) Heatmap of 13 screened chemokines is shown. Quantitative results for chemokines including Ccl2 **(F)**, Ccl4 **(G)**, and Ccl5 **(H)** are shown. **(I-L)** Pancreata from the KC and KC/Pd mice with the same treatment as described in panels **A-H** were harvested and processed for measurements of a panel of 13 cytokines using BioLegend’s LEGENDplex bead-based immunoassays (n=3-5 per group). For GW diet treatment, the pancreatic tissue protein lysates used for **I-J** are the same as used for **A-D**; and for HFD treatment, the pancreatic lysate samples used for **K-L** are same as used for **E-H**. **(I and K)** Heatmaps of 13 screened cytokines from 3 representative mice per group (**I** for GW diet and **K** for HFD) are shown. **(J and L)** Quantitative results of Il6 production for the indicated mouse groups (**J** for GW diet and **L** for HFD). Data are mean ± SEM; \**P* <.05, \*\*\**P* <.001 compared to KC mice with the same diet treatment. **(M)** Heatmap of human pancreatic chemokines and cytokines’ gene expression in normal and pancreatic ductal adenocarcinoma (PDAC), derived from Oncomine Badea database. Enlarged symbols show genes (e.g., Ccl2, Ccl4-5, and Il6) that were in agreement with mouse findings by chemokine and cytokine panel screenings and related to MDSCs’ mobilization and recruitment. Data are mean ± SEM; \**P* <.05, \*\**P* <.001, \*\*\*\**P* <.0001 compared to normal pancreas tissues.

The cytokine panel screening results revealed increases in 7 of 13 inflammatory cytokines (Csf2, Il1a, Il1b, Ifng, Il-6,Tnfa and Mcp1/Ccl2), starting at 3 days of the treatment and with further increases in all 7 cytokines at 9 days, in KC/Pd mice on the GW diet, compared to KC mice on the GW diet or compared to KC/Pd mice on the control diet (0 days of GW) (**Fig. 6I and J, sFig. 5G and H**). In addition, we found that 9 days on the GW diet decreased production of Il23 and Il27 in pancreata of both KC and KC/Pd mice, and of Il10 in pancreata of KC/Pd mice, compared to the same mouse models on the control diet (**Fig. 6I**). Also, at 9 days of the treatment, we saw decreases of Il23, Il27 and Il10 in pancreata of KC/Pd mice on the GW diet, compared to KC mice on the GW diet (**Fig. 6I**). Similarly, we saw increases in Il6, Tnfa, and Mcp1/Ccl2 in KC/Pd mice on a high-fat diet for 12 weeks, compared to either KC mice on a high fat diet or compared to KC/pd mice on a control diet **(Fig. 6K and L, sFig. 5I)**. However, the high fat-diet had no effect on production of Il23, Il27 or Il10 in KC or KC/Pd mice, compared to the same mouse models on the control diet (**Fig. 6I)**.

To evaluate the clinical relevance of our mouse findings, using Oncomine public database analyses, we found that PPARD mRNA expression levels were upregulated in human PDACs, compared to adjacent normal pancreas tissues in the Badea pancreas set, and this PPARD upregulation was associated with significant upregulation of several chemokines such as CCL2, CCL4-5, CXCL5 and cytokines such as IL6 (**Fig. 6M),** which supports our mouse findings.

### Ppard hyperactivation directly upregulates multiple chemokines and cytokines in pancreatic epithelial cells that recruit immunosuppressive cells into pancreata in KC mice

Ppard functions as a transcriptional factor and was overexpressed in the pancreatic epithelial cells that had *Kras^G12D^* mutation in KC/Pd mice. We thus investigated whether Ppard hyperactivation directly upregulated chemokines and cytokines in pancreatic epithelial cells rather than pancreatic stromal cells during the early stage of pancreatic tumorigenesis in KC/Pd mice. In this study, we performed pancreatic epithelial cell tracing by breeding KC/Pd or KC mice with CAG-LSL-tdTomato mice to generate KC/Pd-tdTomato (KC/tdPd) or KC-tdTomato mice (KC/td) mice, in which *Kras^G12D^* mutation, Ppard overexpression and tdTomato red fluorescence protein (RFP) expression or Kras^G12D^ mutation and tdTomato RFP expression, respectively, were directed into Pdx1^+^ pancreatic epithelial cells (**Fig. 7A**). KC/td and KC/tdPd mice at 18 weeks, were fed a GW diet or control diet for 3 days and then sacrificed. We found that the tdTomato RFP–sorted enzymatically digested pancreatic epithelial cells via flow cytometry from KC/td/Pd mice had significantly higher mRNA expression levels of Ccl2, Ccl4, Ccl5, Cxcl5, Il6 than those from sex- and age-matched KC/td littermates did, and these increases were augmented by a GW diet in KC/tdPd mice (**Fig. 7B-D**). These results demonstrated that Ppard hyperactivation directly upregulated a complex mix of multiple chemokines (e.g., Ccl2, Ccl4-5,Cxcl5) and cytokines (e.g.,Il6) in pancreatic epithelial cells, which are known to expand and recruit MDSCs and macrophages into pancreata ^33–35^.

**Figure 7.**
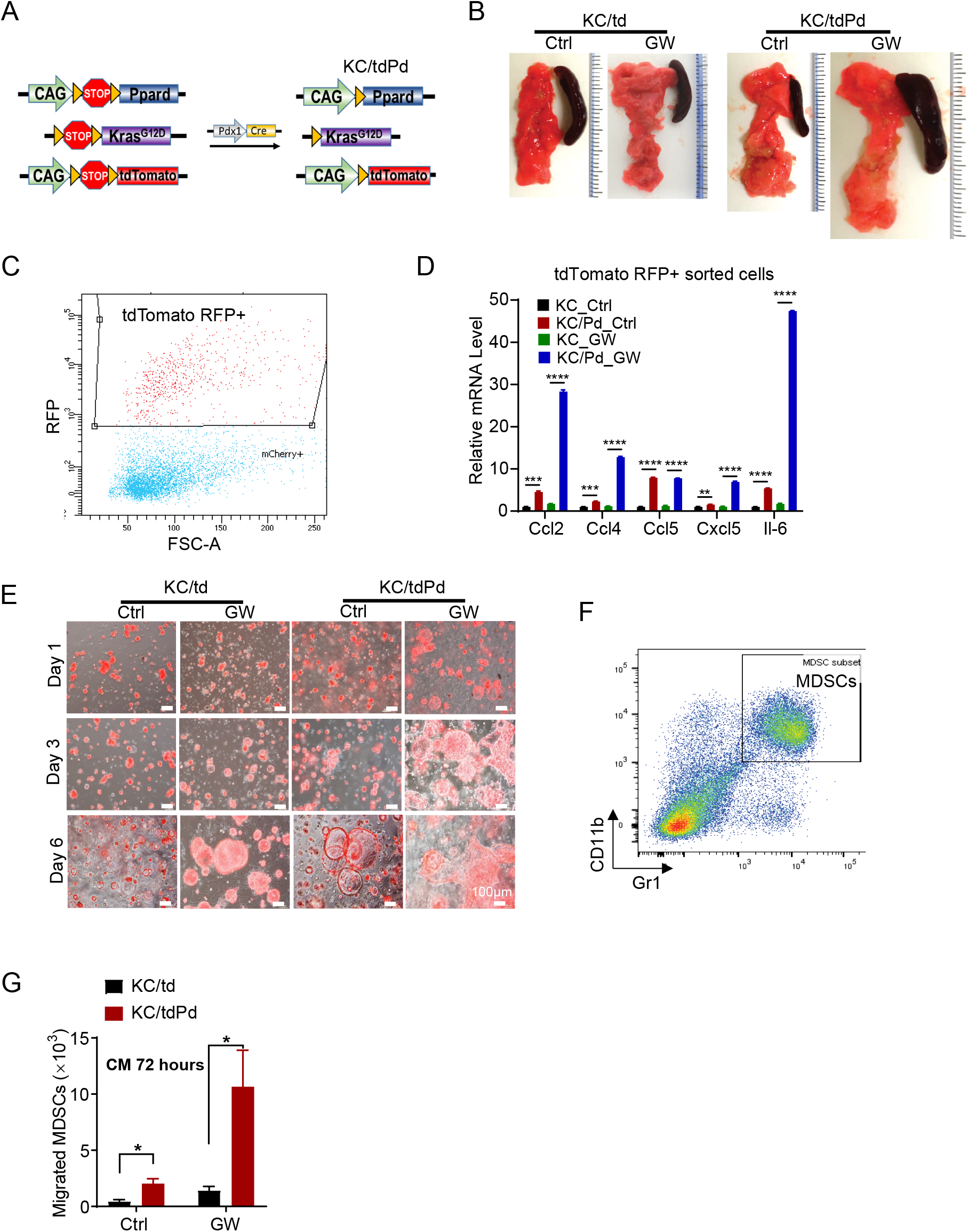
Ppard hyperactivation directly increases production of multiple chemokines and cytokines in pancreatic epithelial cells to chemoattract MDSCs and macrophages into pancreata in KC mice. **(A)** Schematic diagram of the generation of KC/tdPd mice. **(B-D)** KC/td and KC/tdPd mice at 18 weeks old were fed a GW (50 mg/kg) or control (Ctrl) diet for 3 days and then euthanized. **(B)** Representative photos of pancreata for the indicated mouse group. **(C and D)** Pancreata from the indicated mouse groups were enzymatically digested using collagenase IV and DNase I as described in the Methods, sorted by flow cytometry with tdTomato RFP, and harvested for further analyses. **(C)** Representative flow cytometry images of sorted tdTomato RFP^+^ pancreatic epithelial cells. **(D)** The sorted tdTomato RFP^+^ pancreatic epithelial cells from the indicated mouse groups were measured for mRNA expression levels of Ccl2, Ccl4-5, Ccl17, Cxcl5, Tnfa and Il6 by qRT-PCR. The quantitative results are shown. Data are mean ± SEM; \*\*\**P* <.001 and \*\*\*\**P* <.0001. **(E)** KC/td and KC/tdPd mice at 6-8 weeks were fed the GW (50 mg/kg) or control (Ctrl) diet for 3 days and then euthanized. The pancreata were enzymatically digested as described in panels **C and D**, and the digested pancreatic epithelial cells were harvested for 3D organoid culture. GW501516 (1 μM) was added into culture media for the mice fed the GW diet. The organoid images were photographed at 1, 3, and 7 h of culture. Shown are representative merged images with bright-field light and tdTomato-RFP of pancreatic organoids at the indicated time points of primary organoid culture. **(F, G)** MDSCs’ migration towards conditioned medium (CM) harvested from primary pancreatic organoid culture. **(F)** MDSCs (Cd11b^+^Gr1^+^) were isolated from spleens of C57BL/6 wild-type mice by flow cytometry. Representative flow cytometry images of sorted MDSCs are shown. **(G)** Migration assay was performed using 3-μm trans-well inserts with added isolated MDSCs (1×10^5^/well) that were incubated with the harvested CM from the organoid culture at 3 days from panel **F**. After 4 h of incubation, MDSCs that had migrated to the bottom chamber were counted. Quantitative results for migrated MDSCs for the indicated mouse groups are shown (n=3 per group). Data are mean ± SEM; \**P* <.05.

Furthermore, we performed 3D culture of primary pancreatic cell organoids derived from enzymatically digested pancreatic epithelial cells of KC/td and KC/tdPd littermates fed a GW diet or a control diet for 3 days. GW (1 μM) was added in 3D culture media of organoids from mouse groups fed a GW diet. We found that GW treatment increased organoid regenerations as evidenced by increasing organoid sizes in KC/td mice; and this effect of GW was augmented in organoids derived from KC/tdPd, which had a more dedifferentiated appearance (**Fig. 7E**). Conditioned media from 3D culture of GW-treated KC/td organoids at 3 days of culture significantly increased migration of MDSCs (CD11b^+^Gr1^+^), which were isolated from spleens of wild-type C57BL/6 mice (**Fig. 7F**), compared to the conditioned media from 3D culture of control-treated KC/td organoids at the same time point and this effect of GW was strongly augmented in conditioned media from GW-treated KC/tdPd organoids with the same 3D culture (**Fig. 7G**). These results further support Ppard hyperactivation increases the recruitment of MDSCs and thus promoted pancreatic tumorigenesis in KC mice.

## DISCUSSION

In the present study, we found: 1) PPARD expression is upregulated in precancerous human PanINs in humans and KC mice. 2) Ppard activation, by its natural ligands such as fatty acids in a high-fat diet or the highly specific synthetic ligand GW501516, promoted pancreatic tumorigenesis in and shortened the survival of KC mice, and these effects were augmented by Ppard transgenic overexpression in KC/Pd mice. 3) Ppard genetic KO in pancreatic epithelial cells dramatically inhibited the effect of a high-fat or GW501516 diet on promoting pancreatic tumorigenesis in KC mice. 4) Mechanistically, this Ppard activation in pancreatic epithelial cells increased production of multiple chemokines and cytokines to recruit MDSCs and macrophages into pancreata, leading to the establishment of an immunosuppressive tumor microenvironment and inflammation and subsequently accelerating the progression of pancreatic tumorigenesis. Our findings demonstrate that PPARD plays an essential role in promoting oncogenic *Kras^G12D^*–initiated pancreatic tumorigenesis, especially in the setting of high fat or GW501516 exposures.

It has been reported that PPARD is upregulated in human PDACs ^19, 24^. We found for the first time that PPARD is upregulated even in early stages of human and mouse PanINs. This Ppard upregulation was accompanied by higher levels of mutant *Kras* activation as shown by higher pERK1/2 expression in PanINs in KC mice. Similarly, Ppard was significantly upregulated in mouse pancreatic cancer cells derived from inducible *Kas^G12D^* mice that had been treated with doxycycline to activate oncogenic *Kras^G12D^* ^36^. In addition, Ppard was upregulated by mutant *Kras* in rat intestinal epithelial cells via in-vitro studies ^37^. These findings from others and from our current work indicated that Ppard might be a key downstream target of activated *Kras^G12D^*.

Recent studies have shown that an endogenous activity level of mutant *Kras* can be hyperactivated by pancreatic cancer risk factors, including pancreatitis and an obesogenic high-fat diet challenge, which act as a second hit that promoted PDACs with high penetrance ^8, 12^. However, the molecular mediator linking these risk factors to mutant *Kras* hyperactivation remained unclear. We found Ppard activation by fatty acids in a high-fat diet significantly increased pancreatic tumorigenesis in KC mice, which was augmented by Ppard transgenic overexpression in pancreatic epithelial cells in KC/Pd mice. The high-fat diet also increased Ppard expression in both KC and KC/Pd mice. Furthermore, we found that Ppard activation by its specific synthetic ligand GW501516 significantly shortened the survival of KC mice because of the accelerated pancreatic tumorigenesis. Moreover, when Ppard was hyperactivated by GW501516 and Ppard overexpression in pancreatic epithelial cells, pancreatic tumorigenesis was drastically accelerated, leading to quick PDAC development within 9 days and mouse survivor of less than 30 days after GW501516 treatment. The drastic acceleration of *Kras^G12D^*–initiated pancreatic tumorigenesis by Ppard activation demonstrated the essential functions of Ppard hyperactivation in oncogenic *Kras^G12D^*–initiated pancreatic tumorigenesis. These findings are in agreement with prior reports that high-fat diets promote oncogenic *Kras*–initiated pancreatic tumorigenesis ^8^ but also demonstrate for the first time the mechanistic contribution of Ppard hyperactivation to the promotion of oncogenic *Kras*–initiated pancreatic tumorigenesis. Our findings that in a complementary Ppard genetic KO mouse model, the genetic KO of Ppard expression in pancreatic epithelial cells significantly inhibited the pancreatic tumorigenesis promoted by the high-fat or GW501516 diet in KC mice further illustrated the essential and mechanistic role of Ppard in this process. Therefore, Ppard activation by intake of Ppard ligands whether synthetic (e.g., Cardarine, the commercial name for GW501516) or natural (e.g., high levels of fatty acids), can increase the risk of PDAC to humans whose pancreata harbor *Kras* mutations, which are common events along with aging. As a corollary, the development of specific PPARD antagonists to inhibit PPARD activity and function is a potential strategy for PDAC prevention and therapy.

We observed that Ppard hyperactivation induced an immunosuppressive microenvironment that promoted pancreatic carcinogenesis through the recruitment of MDSCs and macrophages. Tumor immunosuppression is emerging as a major pro-tumorigenic mechanisms ^38, 39^. MDSCs are diverse cells that are generated by the pathologic activation of monocytes and immature neutrophils under various disease states (e.g., obesity and cancer) to suppress T cell functions and thus support the disease process ^40^. MDSCs have two major subtypes: PMN-MDSCs and M-MDSCs, which suppress the immune response in antigen-specific and non-specific manners, respectively ^41^. MDSCs expand during tumorigenesis to promote tumor immunosuppression and subsequently tumorigenesis, angiogenesis, and metastasis ^42^. MDSCs subsequently modulate various innate and adaptive immune response cells to create an immunosuppressive microenvironment ^38^. Infiltrating-macrophages in tumors as referred to tumor-associated macrophages (TAMs), are the key components of tumor microenvironment that influences tumor progression ^43^. TAMs can be polarized into two different phenotypes: M1-TAMs act in tumor-inhibiting manner, and M2-TAMs act in tumor-promoting manner. MDSCs and M2-TAMs also strongly promote PDAC development, as demonstrated in various preclinical models of oncogenic *Kras*-initiated pancreatic tumorigenesis ^33, 34, 44^. Promotion of oncogenic *Kras*–initiated pancreatic tumorigenesis by high-fat diets is also associated with tumor immunosuppressive microenvironment ^8, 9, 45^. Our own current immunophenotyping studies of the pancreatic microenvironment showed that Ppard hyperactivation by a high-fat or GW diet markedly increased immunosuppressive cells such as MDSCs and macrophages (e.g., M2-TAMs), and markedly decreased CD8^+^ effector T cells, in pancreata from KC/Pd mice. MDSCs and M2-TAMs have emerged as important factors for the progression of mutant *Kras*-initiated pancreatic tumorigenesis by facilitating tumor escape of immune surveillance ^33, 44^.

Pancreatic ductal cells undergoing malignant transformation are able to mobilize and recruit MDSCs to pancreata to create tumor immunosuppression by producing various chemokines and cytokines (e.g., Ccl2, Ccl4-5, Cxcl5, Il6) ^33, 35, 46^. Indeed, our comparative RNA-seq transcriptome profiling studies revealed that inflammatory response signaling pathways were among the top pathways that promoted pancreatic tumorigenesis with Ppard hyperactivation. When we screened the pancreata from the same mouse models that were used for immunophenotyping studies to identify the chemokines and cytokines that triggered this immunosuppressive microenvironment, we found that Ppard hyperactivation via on the GW diet for 3 days or on a high-fat diet for 12 weeks, increased various chemokines (e.g., Ccl2, Ccl4-5, Cxcl5) and cytokines (e.g., Il6), recruiting MDSCs and macrophages into the pancreas ^33, 34, 46–48^. To assess the clinical relevance of these findings, we searched the Oncomine database, and found that PPARD is upregulated in human PDACs, which was associated with upregulation of these various pro-tumorigenic chemokines (e.g., CCL2, CCL4-5, and CXCL5) and cytokines (e.g., IL6) in agreement with our mouse findings. These data strongly support the notion that Ppard hyperactivation by a high fat diet accelerates pancreatic tumorigenesis via increasing chemokine-mediated recruitment of immunosuppressive cells into pancreata in our mouse models.

We found that Ppard hyperactivation to increase production of these chemokines and cytokines occurs in pancreatic epithelial cells directly. Our cell lineage tracing studies confirmed that Ppard-hyperactivated Pdx1^+^ pancreatic epithelial cells, sorted by flow cytometry with tdTomato RFP, produced significantly higher levels of the chemokines and cytokines (e.g., Ccl2, Ccl4-5, Cxcl5, Il6) identified by our screening, which are known to recruit immunosuppressive cells such as MDSCs and M2-TAMs in KC mice. On the other hand, normal pancreatic epithelial cells from C57BL/6 wild-type mice typically had low or no expression of these chemokines and cytokines. Our results clearly demonstrated the essential role of PPARD in promoting pancreatic tumorigenesis by affecting multiple facets of tumor progression.

In summary, Ppard plays an essential role in promoting mutant *Kras*–initiated pancreatic tumorigenesis, especially in the setting of a high-fat diet or GW501516 treatment. Our findings provide a strong rationale for targeting PPARD as a novel interventive strategy to prevent and suppress development and progression of PDAC, especially for patients with early stage of pancreatic tumors with upregulated PPARD expression.

## Supporting information

Supplemental Figures 1-5 with legends

## STAR * METHODS

Detailed methods are provided in the online version of this paper and include the following:

- KEY RESOURCES TABLE
- RESOURCE AVAILABILITY

- Lead contact
- Materials availability
- Data and code availability
- EXPERIMENTAL MODEL AND SUBJECT DETAILS

- Human tissue samples
- Animals
- METHOD DETAILS

- Generation of experimental mice
- Mouse treatment with high-fat or GW501516 diet and pancreatic tumorigenesis evaluation
- Mouse survival experiments
- Histology
- Immunohistochemistry and immunofluorescence staining
- RNAscope in situ hybridization
- RNA extraction and quantitative real-time RT-PCR
- RNA sequencing
- Protein lysate preparation and Western blotting
- Flow cytometry and pancreatic acinar cell sorting
- Quantification of a panel of LEGENDplex mouse proinflammatory chemokines or cytokines
- Immune cell profiling by flow cytometry
- Primary pancreatic organoid culture
- QUANTIFICATION AND STATISTICAL ANALYSIS

## SUPPLEMENTAL INFORMATION

Supplemental Information can be found online.

## ACKNOWLEDGMENTS

This work was supported by the National Cancer Institute (R01-CA142969, R01-CA195686, R01-CA206539 to I.S., and R03 CA235106 to X. Zuo), the Cancer Prevention and Research Institute of Texas (RP150195 and RP140224 to I.S.) and MD Anderson faculty incentive fund (to I.S.). This study made use of the MD Anderson Cancer Center Genetically Engineered Mouse Facility, Functional Genomics Core, Flow Cytometry and Cellular Imaging Facility, the Next Generation Sequencing Core and Research Animal Support Facility—Smithville Laboratory Animal Genetic Services, supported by Cancer Center Support Grant P30CA016672. The Next Generation Sequencing Core was also supported by CPRIT Core Facility Support Grant RP120348. We thank the Dr. Craig D. Logsdon lab at The University of Texas MD Anderson Cancer Center for providing the CAG-LSL-*Kras^G12V^* plasmid for generation of CAG-LSL-Ppard DNA fragments for pronuclear injection as described in Animals section.

We thank Ms. Sarah J Bronson at the Department of Scientific Publications at MD Anderson Cancer Center for editing the manuscript.

## AUTHOR CONTRIBUTIONS

I.S. and X. Zuo conceived the study and designed the experiments. Y.L., Y.D., X.Zuo, L.A.V., and F.L. performed various portions of animal experiments and the in vitro experiments. D.W. contributed to the experimental designs and provided technical support for animal and the in vitro experiments. M.J.M. performed three-dimensional organoid culture. Y.L., X. Zuo, and J.K.C. performed flow cytometry analyses for pancreata-infiltrating immune cell profiles, and Y.L. evaluated the results. H.W. provided human pancreas tissue sections and edited the manuscript. H.W., Y.D., X. Zuo., and Y.L. performed pathologic evaluations of mouse and human tissue samples. D.L. coordinated with the Next Generation Sequencing Core at MD Anderson for performing RNA sequencing transcriptome profiling assays. X. Zheng, Y.L., and J.W. evaluated RNA sequencing results and performed gene set enrichment analyses. B.J. constructed the CAG-Loxp-GFP-Loxp-*KRAS^G12V^* plasmid. I.S., X. Zuo, Y.L., and Y.D. analyzed the data. I.S., X. Zuo, and Y.L. wrote the manuscript. D.W., D.L., B.J., and J.C.Y. provided conceptual feedback for the manuscript.

## STAR * METHODS

### KEY RESOURCES TABLE

| REAGENT or RESOURCE | SOURCE | IDENTIFIER |
| --- | --- | --- |
| Antibodies |  |  |
| Mouse anti-α-SMA (IF) | Sigma-Aldrich | A5228 |
| Rabbit phospho-ERK1/2 (IHC) | Cell Signaling Technology | 4370S |
| Rabbit polyclonal anti-Ppard (WB) | Abcam | Ab8937 |
| Rabbit polyclonal anti-human PPARD (IHC) | Aviva Systems Biology | ARP38765_T100 |
| Rat monoclonal anti-CK19 (IF) | Developmental Studies Hybridoma Bank | TROMA-III-c |
| Rabbit polyclonal anti-α-amylase (IF) | Sigma-Aldrich | A8273 |
| PE monoclonal anti-mouse I-A/I-E (F) | BioLegend | 107607 |
| PerCP/Cy5.5 monoclonal anti-mouse CD45 (F) | BioLegend | 103131 |
| Rat Brilliant Violet 785 monoclonal anti-mouse CD3ε (F) | BioLegend | 100355 |
| Rat Alexa Fluor 700 monoclonal anti-mouse/human CD45R/B220 (F) | BioLegend | 103231 |
| Rat Brilliant Violet 510 monoclonal anti-mouse Ly-6G/Ly-6C (Gr1) (F) | BioLegend | 108438 |
| Rat PE/Cy7 monoclonal anti-mouse/human CD11b (F) | BioLegend | 101215 |
| Rat Brilliant Violet 421 monoclonal anti-mouse F4/80 (F) | BioLegend | 123131 |
| Rat Brilliant Violet 510 monoclonal anti-mouse Ly6C (F) | BioLegend | 128033 |
| Rat APC/Fire 750 monoclonal anti-mouse Ly6G (F) | BioLegend | 127651 |
| Rat BUV737 monoclonal anti-mouse CD8α (F) | BD Biosciences | 564297 |
| Rat FITC monoclonal anti-mouse CD4 (F) | BioLegend | 100405 |
| TruStain FcX (anti-mouse CD16/32) antibody (F) | BioLegend | 101319 |
| Biotinylated goat anti-rabbit IgG antibody | Vector Laboratories | BA-1000 |
| Goat anti-rat IgG (H+L) secondary antibody, Alexa Fluor 594 conjugated | Thermo Fisher Scientific | A-11007 |
| Goat anti-rabbit IgG (H+L) secondary antibody, Alexa Fluor 488 conjugated | Thermo Fisher Scientific | A-11034 |
| Goat anti-rabbit IgG (H+L) secondary antibody, Alexa Fluor 594 conjugated | Thermo Fisher Scientific | A-11012 |
| Biological Samples |  |  |
| Human paraffin-embedded pancreas tissue array | US Biomax | BIC14011a |
| Chemicals, Peptides, and Recombinant Proteins |  |  |
| 10% buffered formalin | Thermo Fisher Scientific | 23-245-685 |
| 4% paraformaldehyde solution | Thermo Fisher Scientific | AAJ19943K2 |
| Advanced Dulbecco modified Eagle medium (DMEM)/Ham F-12 medium | Thermo Fisher Scientific | 12634010 |
| RPMI 1640 medium | Thermo Fisher Scientific | 11875119 |
| McCoy 5A (modified) medium | Thermo Fisher Scientific | 16600082 |
| Antigen unmasking solution | Vector Laboratories | H3300 |
| Collagenase, type IV | Sigma-Aldrich | C5138 |
| DMEM | Thermo Fisher Scientific | 11995073 |
| Dimethyl sulfoxide | Sigma-Aldrich | D2650 |
| ECL Western blotting substrate | Thermo Fisher Scientific | PI32209 |
| Eosin-Y 7111 | Thermo Fisher Scientific | 22-050-110 |
| Ethylenediaminetetraacetic acid (EDTA) | Sigma-Aldrich | E9884 |
| Fetal bovine serum | Atlanta Biologicals | S12450H |
| GlutaMAX (100x) | Thermo Fisher Scientific | 35050061 |
| Normal goat serum | Cell Signaling Technology | 5425S |
| Hexadimethrine bromide (polybrene) | Santa Cruz Biotechnology | SC-134220 |
| Hydrogen peroxide | Sigma-Aldrich | H3410 |
| Matrigel | Corning | 356231 |
| Mayer's hematoxylin | Agilent Dako | S330930-2 |
| Nonidet P-40 | Sigma-Aldrich | 74385 |
| *N*-acetyl-L-cysteine | Sigma-Aldrich | A7250 |
| Penicillin-streptomycin-glutamine (100x) | Thermo Fisher Scientific | 10378016 |
| Phenylmethylsulfonyl fluoride | Sigma-Aldrich | P7626 |
| Recombinant murine Wnt-3a | EMD Millipore | GF154 |
| TRIzol | Thermo Fisher Scientific | 15596018 |
| Trypsin-EDTA | Thermo Fisher Scientific | 25300062 |
| Recombinant human Wnt-3a protein | R&D Systems | 5036-WN-010 |
| cOmplete protease inhibitor cocktail | Roche | 11697498001 |
| Ponceau S solution | Sigma-Aldrich | P7170 |
| Percoll density gradient solution | Fisher Sci (GE) | 45-001-747 |
| Collagenase (type 8) | Sigma-Aldrich | C-2139 |
| DNase I | Sigma-Aldrich | DN25 |
| DL-Dithiothreitol | Sigma-Aldrich | D9779 |
| RBC lysis buffer (10X) | BioLegend | 420301 |
| Cell staining buffer | BioLegend | 420201 |
| True-Nuclear Transcription Factor Buffer Set | BioLegend | 424401 |
| Critical Commercial Assays |  |  |
| iScript cDNA Synthesis Kit | Bio-Rad | 1708891BUN |
| FastStart Universal Probe Master (Rox) | Roche | 4914058001 |
| ProLong Gold Antifade Mountant with DAPI | Thermo Fisher Scientific | P36935 |
| LEGENDplex Mouse Proinflammatory Chemokine Panel (13-plex) | BioLegend | 740007 |
| LEGENDplex Mouse Inflammation Panel (13-plex) | BioLegend | 740150 |
| NovaUltra Alcian blue stain kit | IHCWORLD | IW-3000 |
| NovaUltra Sirius red stain kit | IHCWORLD | IW-3012 |
| Zombie UV Fixable Viability Kit | BioLegend | 423107 |
| ArC Amine Reactive Compensation Bead kit | Thermo Fisher Scientific | A10628 |
| UltraComp Beads for Ab single stains | Thermo Fisher Scientific | 01-2222-41 |
| Precision Count Beads | BioLegend | 424902 |
| RNAscope 2.5 HD Reagent Kit-RED | Biotechne ACDBio | 322350 |
| Experimental Models: Organisms/Strains |  |  |
| Mouse: B6.FVB-Tg(Pdx1-cre)6Tuv/J | Jackson Laboratory | 014647 |
| Mouse: C57BL/6J (B6) | Jackson Laboratory | 664 |
| Mouse: B6.Cg-Gt(ROSA)26Sor<sup>tm14(CtdTomato)Hze</sup>/J | Jackson Laboratory | 007914 |
| Mouse: B6.129S4-Kras<sup>tm4Tyj</sup>/J | Jackson Laboratory | 008179 |
| Mouse: PPAR-d-flox | Donated by Dr. Ronald Evans | N/A |
| Mouse: CAG-LSL-PPAR-d<sup>OE</sup> | Generated by our lab | N/A |
| Mouse: FVB/N | Charles River Laboratories | 207 |
| Oligonucleotides |  |  |
| Mouse Ccl2 (FAM/MGB) | Thermo Fisher Scientific | Mm00441242_m1 |
| Mouse Ccl17 (FAM/MGB) | Thermo Fisher Scientific | Mm01244826_g1 |
| Mouse Tnf (FAM/MGB) | Thermo Fisher Scientific | Mm00443258_m1 |
| Mouse Il6 (FAM/MGB) | Thermo Fisher Scientific | Mm00446190_m1 |
| Mouse Ywhaz (FAM/MGB) | Thermo Fisher Scientific | Mm01158417_g1 |
| Mouse Ppard (FAM/ZEN/IBFQ) | Integrated DNA Technologies | Mm.PT.58.6994542 |
| Mouse Ccl3 (FAM/ZEN/IBFQ) | Integrated DNA Technologies | Mm.PT.58.29283216 |
| Mouse Ccl4 (FAM/ZEN/IBFQ) | Integrated DNA Technologies | Mm.PT.58.5219433 |
| Mouse Ccl5 (FAM/ZEN/IBFQ) | Integrated DNA Technologies | Mm.PT.58.43548565 |
| Mouse Csf2 (FAM/ZEN/IBFQ) | Integrated DNA Technologies | Mm.PT.58.9186111 |
| Mouse Cxcl5 (FAM/ZEN/IBFQ) | Integrated DNA Technologies | Mm.PT.58.29518961.g |
| RNAscope Probe-Mm-Ppard | Biotechne ACDBio | 504451 |
| RNAscope Probe-Hs-PPARD | Biotechne ACDBio | 546391 |
| Software and Algorithms |  |  |
| Thermo TraceFinder | Thermo Fisher Scientific | N/A |
| GraphPad Prism 7 | GraphPad Software Inc. | SCR_002798 |
| LEGENDplex v8.0 | BioLegend | N/A |
| FlowJo v10 | BD FlowJo | N/A |
| RStudio | https://rstudio.com/ | N/A |
| Rv4.0.2 | https://cran.r-project.org/mirrors.html | N/A |
| SAS software 9.4 | SAS Institute | N/A |
| Adobe Illustrator CC 2017 | Adobe Inc. | SCR_010279 |

### RESOURCE AVAILABILITY

#### Lead Contact

Further information and requests for resources and reagents should be directed to and will be fulfilled by the lead contact, Imad Shureiqi.

#### Materials Availability

Unique reagents and materials generated in this study are available after completion of an institutionally required Material Transfer Agreement.

#### Data and Code Availability

The RNA-seq dataset generated during this study is under processing to be deposited to the NCBI Gene Expression Omnibus public database.

### EXPERIMENTAL MODEL AND SUBJECT DETAILS

#### Human pancreas tissue sections and human pancreas tissue microarrays

A set of pancreatic tissue microarray including paired normal, PanINs and PDAC were purchased from US Biomax (#BIC14011a).

Human pancreatic tissue samples were collected from patients undergoing endoscopic ultrasound guided biopsies of pancreatic tumors at MD Anderson Cancer Center. A written informed consent was obtained from each patient. The current study using these tissue samples was conducted in accordance with the recognized ethical guidelines (Declaration of Helsinki, CIOMS, Belmont Report, and U.S. Common Rule) and approved by The Institutional Review Board of MD Anderson. De-identified sections from freshly formalin-fixed pancreatic tissues were used for performing hematoxylin and eosin (H&E) staining and PPARD RNAscope in situ hybridization assays. Histologic evaluation of the H&E staining was performed by an experienced pancreatic oncologic pathologist (H.W.).

#### Animals

All animal experiments were conducted in accordance with the Guide for the Care and Use of Laboratory Animals and were approved by Institutional Animal Care and Use Committee of MD Anderson. *Ppard* conditional overexpression transgenic mice (CAG-LSL-*Ppard*) were generated by pronuclear injection of sequence-confirmed purified DNA fragments of CAG-LoxP-GFP-LoxP-*Ppard* at the Genetically Engineered Mouse Facility at MD Anderson, in which a mouse *Ppard* full-length cDNA was subcloned into HindIII/EcoRV sites of pCAGGS-LoxP-GFP-LoxP-*Kras^G12V^* vector to replace the *Kras^G12V^*. The resultant vector was digested with PmeI to remove vector sequences, and the DNA fragment of CAG-LoxP-GFP-LoxP-*Ppard* driven by a CAG promoter (containing the chicken β-actin promoter, cytomegalovirus IE enhancer, and β-actin intron) was purified for this pronuclear injection. The same vector backbone has been widely used in generating the transgenic mice except with *Ppard* replacement of NF-κB ^49^, IKK2 ^50^, or KLF4 ^51^. For genotyping of CAG-LSL-*Ppard* transgenic mice, mouse tail genomic DNA samples were used in qPCR analyses using a mouse *Ppard* PCR primers and probe set (Life Technologies, Assay ID: Mm00803184_m1). *Ppard* conditional KO mice, in which *Ppard* exon 4 is flanked with *loxP* sites (designated as Ppard-flox mice), were a gift from Dr. Ronald Evans ^52^. Pdx1-*Cre* (#014647), LSL-*Kras*^G12D^ (#008179), and B6.Cg-*Gt(ROSA)26Sor^tm14(CAG-tdTomato)Hze^/J* (LSL-tdTomato, #007914) mice were purchased from the Jackson Laboratory.

### METHOD DETAILS

#### Generation of experimental mice

1. *Pdx1-Cre* mice were bred with LSL-*Kras*^G12D^ or with LSL-*Kras*^G12D^; CAG-LSL-*Ppard* mice to generate *Pdx1-Cre;* LSL-*Kras*^G12D^ mice (called KC mice); or *Pdx1-Cre;* LSL-*Kras*^G12D^; CAG-LSL-*Ppard* mice (called KC/Pd mice), in which *Kras^G12D^* mutation or both *Kras*^G12D^ mutation and *Ppard* overexpression, respectively, were induced specifically in Pdx1-expressing pancreatic epithelial cells.
2. *Pdx1-Cre; Ppard*-flox^(+/-)^ mice were bred with *LSL-Kras^G12D^; Ppard*-flox^(+/-)^ mice to generate *Pdx1-Cre;* LSL-*Kras*^G12D^; *Ppard*-flox^(-/-)^ mice (called KC/PdKO mice), in which *Kras^G12D^* mutation and *Ppard* genetic deletion/KO were induced specifically in Pdx1-expressing pancreatic epithelial cells.
3. LSL-tdTomato mice were bred with KC or KC/Pd mice to generate *Pdx1-Cre; LSL-Kras^G12D^;* LSL-tdTomato or *Pdx1-Cre; LSL-Kras^G12D^;* LSL-tdTomato; LSL-*Ppard*^OE^ mice, in which Pdx1-expressing epithelial cells were marked with tdTomato fluorescence protein while *Kras^G12D^* mutation (KC/td) or *Kras^G12D^* mutation and Ppard overexpression (KC/tdPd) were directed into these cells.

#### Mouse treatment with high-fat or GW501516 diet and pancreatic tumorigenesis evaluation

<u>*1)*</u> KC or KC/Pd mice at age 6-8 weeks were fed either a high-fat diet (60% kcal from fat, Envigo) or a control diet (10% kcal from fat, Envigo) for 12 weeks (n=4-10 mice per group). <u>*2)*</u> KC or KC/Pd mice at age 6-8 weeks were fed a diet containing 50 mg/kg GW501516 (Envigo) for 0 days (control diet: the same diet except without GW501516, for 9 days), 3 days (control diet for 6 days and GW diet for 3 days), or 9 days (GW diet for 9 days) (n=5-8 mice per group). The chemical GW501516 was synthesized by the Translational Chemistry Core Facility at MD Anderson Cancer Center, and its authenticity was confirmed by liquid chromatography–mass spectrometry using standard GW501516 (#SML1491, Sigma-Aldrich). <u>*3)*</u> KC or KC/PdKO mice at age 6-8 weeks were fed either a high-fat diet (60% kcal from fat, Envigo) or a control diet (10% kcal from fat, Envigo) for 26 weeks (n=5-10 mice per group). <u>*4)*</u> KC or KC/PdKO mice at age 6-8 weeks were fed a diet containing 50 mg/kg GW501516 or control diet (Envigo) for 13 weeks (n=5-8 mice per group). The mice were euthanized after the completion of the treatment, and the pancreas from each mouse was removed and grossly inspected for tumor formation. Half of the pancreas from each mouse was harvested for RNA and protein analyses, and the other half was put in 10% neutral formalin to fix overnight. The formalin-fixed tissues were placed in cassettes, dehydrated in alcohol gradients, and then embedded in paraffin for further sectioning analysis (H&E, IHC, or IF staining). H&E-stained sections were classified and graded with the support of an experienced pancreatic pathologist.

#### Mouse survival experiments

The survival experiments were performed on KC or KC/Pd mice at age 6-8 weeks fed a GW501516 (50 mg/kg) or a control diet. The mice were followed until they required euthanasia on the basis of one of the following preset criteria with the support of experienced veterinary technologists from our animal facility: (1) moribundity, dyspnea, anemia, hunched posture, rough coat or/and (2) weight loss of more than 20%.

#### Histology

Formalin-fixed samples were embedded in paraffin and sectioned onto slides at 5-μm thick and then stained with H&E. Histologic assessment was performed with the support of an experienced pancreatic pathologist. PanINs (1-3) and PDAC lesions in the mice were evaluated by an experienced pathologist based on criteria described at http://pathology.jhu.edu/pc/professionals/DuctLesions.php. All foci of PanINs (1-3) or PDAC lesion areas were counted, and frequencies of PanINs (1-3) and PDAC were then calculated as the percentage of the pancreatic neoplastic area out of the whole pancreas area.

#### IHC and IF staining

IHC and IF staining was performed as described previously ^53^. Tissue sections 5 μm thick were deparaffinized and rehydrated. Antigen retrieval was then performed by immersion of slides into Antigen Unmasking Solution (#H3300, Vector Laboratories) and heating in a steam chamber for 35 min. For immunohistochemistry staining, slides were treated with 3% H_2_O_2_ and water solution to reduce endogenous peroxidase, incubated with blocking buffer (5% goat serum in TBST) for 30 min, and then incubated with primary antibodies overnight. The following primary antibodies were used: PPARD (for human tissue section: #ARP38765_T100; Aviva Systems Biology) and phospho-ERK1/2 (for both mouse and human tissue sections: #4370S, Cell Signaling Technology). On the second day, the tissue sections were incubated with biotinylated secondary antibodies (VECTASTAIN ABC kit; Vector Laboratories) for 1 h, followed by incubation with avidin-coupled peroxidase (Vector Laboratories) for 30 min. 3,3’-diaminobenzidine (Agilent Dako) was used as the chromogen, and the slides were counterstained with Mayer’s hematoxylin (Agilent Dako). For IF staining, the slides were directly incubated with blocking buffer (1.5% goat serum, 0.3% Triton X-100 in phosphate-buffered saline [PBS]) and then underwent primary incubation overnight. The primary antibodies were: α-SMA (#A5228, Sigma-Aldrich), cytokeratin 19 (TROMA-III-c, Developmental Studies Hybridoma Bank), and α-amylase (#A8273, Sigma-Aldrich) were used. On the second day, the slides were incubated with the Alexa Fluor 488 or Alexa Fluor 594 fluorescence-conjugated secondary antibody at 37°C for 1 h. Slides were then washed and mounted with ProLong Gold Antifade Mountant with DAPI (Thermo Fisher Scientific). IHC scoring was performed as follows: the intensity of staining was categorized as 1 (weak staining), 2 (moderate staining), or 3 (strong staining), and the final score was calculated by multiplying the percentage of positive cells (0-100) and the intensity (0-3) ^54^.

#### RNAscope in situ hybridization and quantifications

RNAscope in situ hybridization was performed according to the manufacture’s manuals (RNAscope 2.5 assay, #322350, Biotechne ACDbio). Briefly, fresh paraffin-embedded pancreatic tissue blocks (blocks embedded within 6 months are required for having integrated mRNA in tissue sections) were sectioned into 5 μm per slide the day before the in situ hybridization experiment. On the next day, the slides were deparaffinized, then treated with 3% H2O2 to block endogenous peroxide at room temperature for 10 min. The slides were immersed in RNAscope specific target retrieval reagent (Biotechne ACDbio) and boiled in a steam chamber for 15 min. Slides were washed by 100% ethanol at room temperature for 3 min, dried at 60°C for 5 min, and then incubated with protease reagent in the humidity control tray in the HybEZ oven (Biotechne ACDbio) at 40°C for 30 min. Next, probe hybridization was performed in the HybEZ oven for 2 h using the following probes: mouse Ppard (#504451, Biotechne ACDbio) and human PPARD (#546391, Biotechne ACDbio). After probe hybridization, the slides were washed, and signal amplification was performed using AMP1-6 reagents (Biotechne ACDbio). The final signals were produced by incubation of slides with Fast RED A and Fast RED B reagents, followed by Mayer’s hematoxylin staining. RNAscope intensity was scored as follows: the staining red dots per cell in each slide were counted under 20/40× bright-field microscopy and then were scored as 0 (no staining or less than 1 dot/10 cells), 1+ (1-3 dots/cell), 2+ (4-10 dots/cell, very few dot clusters), 3+ (>10 dots/cell, and less than 10% positive cells have dot clusters), or 4+ (>10 dots/cell, and more than 10% positive cells have dot clusters). Detailed scoring information can be found at the manufacturer’s guideline (SOP 45006, Biotechne ACDbio).

#### RNA extraction and qRT-PCR

The pancreatic tissues from the euthanized mice were injected with RNAlater solution (Thermo Fisher Scientific) at multiple points using a 30-gauge needle syringe. Then, the tissues were cut into tiny pieces (~1 mm), immersed in RNAlater, and then immediately flash-frozen in liquid nitrogen until the tissues were further processed for RNA analysis. For extraction of total RNA, RNAlater-treated pancreatic tissues were first homogenized in TRIzol by mechanical homogenizer on ice, and then total RNA was isolated according to the TRIzol manufacturer’s instructions. mRNA relative expression levels of mouse Ppard, Ccl2, Ccl4, Ccl5, Cxcl5, Il6, Ccl17 and Tnfα were measured by qRT-PCR as described before ^53^. Briefly, the total RNA was first transcribed to cDNA using the iScript cDNA synthesis kit (Bio-Rad), and then qRT-PCR was performed in the StepOnePlus PCR system (Applied Biosystems) using KAPA Probe Fast qPCR Master Mix (KAPA Biosystems). The mouse Ywhaz gene was used as an endogenous control ^55^.

#### RNA-seq for transcriptome profile analyses

The KC and KC/Pd mice at 6-8 weeks were fed a GW diet (50mg/kg) for 3 days and then euthanized (n=3 per group). Total RNA was isolated from the mouse pancreatic tissues using the RNeasy Mini/Micro Kit (Qiagen) according to the manufacturer’s instructions. Mouse pancreatic tissues were harvested and processed prior to total RNA extraction as described above. The RNA quality was measured by the Agilent Bioanalyzer. The total RNA samples with RNA integrity numbers (RINs) of ≥7 were processed for RNA-seq transcriptome profile analysis in the Next Generation Sequencing Core at the Science Park MD Anderson Cancer Center as described before ^32^. Briefly, the TruSeq RNA sample prep kit v2 (Illumina) was used for constructing Illumina-compatible mRNA libraries and an Illumina HiSeq 2000 instrument was employed for multiplexing and sequencing of the six libraries using the 75-bp paired-end format. After sequencing, the sequencing generated BCL files were converted into.fastq.gz files, and the sample libraries were de-multiplexed using CASAVA 1.8.2 to exclude mismatches. Comparative bioinformatics statistical analyses for the RNA-seq raw data were performed by two experienced experts at the Department of Bioinformatics and Computational Biology at MD Anderson Cancer Center. A gene set enrichment assay for the RNA-seq results was analyzed using the R package “ClusterProfiler” and other associated R packages. The full RNA-seq data are currently under processing to be deposited to the NCBI Gene Expression Omnibus public database.

#### Protein lysate processing and Western blotting

Soon after mice were euthanized, their pancreatic tissues were flash-frozen by liquid nitrogen in sterile tubes. For protein lysate preparation, the frozen tissues were wrapped in liquid nitrogen–chilled, clean foil, ground into powder, and homogenized in protein lysis buffer (1% Nonidet P-40, 20 mM 3-[N-morpholino] propanesulfonic acid [pH 7.0], 2 mM ethylene glycol tetraacetic acid, 5 mM ethylenediaminetetraacetic acid, 30 mM sodium fluoride, 40 mM β-glycerophosphate, 2 mM sodium orthovanadate, 1 mM phenylmethylsulfonyl fluoride [Sigma-Aldrich], and 1× complete protease inhibitor cocktail [Roche Applied Science]). A concentration of 10%-12% sodium dodecyl sulfate polyacrylamide gel was used for electrophoresis. The protein was transferred to nitrocellulose membrane, blocked with 5% non-fat milk in TBST, and incubated with primary antibodies overnight. The primary antibody used was rabbit polyclonal anti-Ppard (Abcam, #ab8937). Ponceau S–stained total protein in membrane was used as a loading control.

#### Flow cytometry and cell sorting

Mouse pancreatic tissues from the KC/td or KC/tdPd mice fed a GW501516 (50 mg/kg) or control diet for 3 days were rinsed by cold PBS, cut into 1-mm pieces, and incubated with digestion buffer (1 mg/mL collagenase IV [Sigma-Aldrich] and 10 U/mL DNase I [Sigma-Aldrich] in DMEM) at 37°C for 30 min. The digested tissues were pushed through a 70-μm cell strainer, washed by DMEM, and re-suspended in DMEM containing 1% fetal bovine serum and 2 mM EDTA. Digested cells were then sorted by flow cytometry using td-Tomato RFP and harvested in the fresh complete DMEM medium. The cells were centrifuged at low speed of 300 g and subsequently used for further RNA analyses.

#### Quantification of panel of LEGENDplex mouse proinflammatory chemokines or cytokines

The panel profiling of proinflammatory chemokines (BioLegend, #740007) or cytokines (BioLegend, #740150) was performed as described before ^32^. The assay uses the principles of a sandwich enzyme-linked immunosorbent assay to quantify soluble analytes using a flow cytometer, which allows simultaneous quantification of 13 mouse chemokines (Ccl2, Ccl5, Cxcl10, Ccl11, Ccl17, Ccl3, Ccl4, Cxcl9, Ccl20, Cxcl5, Cxcl1, Cxcl13, and Ccl22) or 13 cytokines (Il1α, Il1β, Il6, Il10, Il12 p70, Il17a, Il23, Il27, Mcp1, Ifnb, Ifng, Tnfa, and Gm-Csf). The whole-protein lysates of the pancreatic tissue were processed and harvested as described previously for Western blot, except without adding sodium dodecyl sulfate. The data were analyzed using the LEGENDplex Data Analysis Software. The final results for protein lysates were normalized to their corresponding protein and presented as pg per mg of protein and for sera were presented as pg/ml.

#### Immune cell profiling by flow cytometry

Mouse pancreatic tissues were digested by digestion buffer (1 mg/mL collagenase IV [Sigma-Aldrich] and 10 U/mL DNase I [Sigma-Aldrich] in DMEM) at 37°C for 30 min. The digested tissues were pushed through a 70-μm cell strainer and washed by 40ml DMEM medium without FBS. The digested cells were re-suspended in 37% Percoll, with the same volume of 70% Percoll on the bottom of the tube. After centrifugation at 800g for 20 min with break off, the lymphocytes were isolated from the 37%/70% interface and rinsed by PBS. The cells were first stained by Zombie UV (BioLegend, #423107), then were incubated with a cocktail comprising the following antibodies in cell staining buffer (#420201, BioLegend): anti-CD45 (#103131), anti-CD3ε (#100355), anti-B220 (#103231), anti-Gr1 (#108438), anti-CD11b (#101215), anti-F4/80 (#123131), anti-Ly6C (#128033), anti-Ly6G (#127651), anti-CD4 (#100405), anti-IA/IE (#107607, all from BioLegend), and anti-CD8a (#564297, BD Biosciences). After washing, the stained cells were submitted for multiple-color flow cytometry analysis. The analysis results were further processed using FlowJo software.

#### Primary pancreatic organoid culture

The mouse pancreatic tissues from the KC/td and KC/tdPd mice littermates treated with the GW501516 diet (50 mg/kg, Envigo) or control diet for 3 days (n=5 per group) were digested with digestion buffer (1 mg/mL collagenase IV [Sigma-Aldrich] and 10 U/mL DNase I [Sigma-Aldrich] in DMEM) at 37°C for 30 min, and the pancreatic epithelial cells were harvested, counted, and embedded in Matrigel (Corning). The Matrigel was topped with organoid culture medium comprising advanced DMEM/F-12 supplemented with penicillin-streptomycin, 2 mM GlutaMAX, 10 mM HEPES (all from Thermo Fisher Scientific, Waltham, MA), mouse recombinant Wnt-3a at 100 ng/mL (MilliporeSigma), mouse epidermal growth factor at 50 ng/mL (Invitrogen), mouse noggin at 100 ng/mL (PeproTech), human R-spondin-1 at 1 μg/mL (PeproTech), *N*-acetyl-L-cysteine at 1 mM (Sigma-Aldrich), N-2 1× (Thermo Fisher Scientific), B-27 1× (Thermo Fisher Scientific), and Y-27632 at 10 μM (Fisher Scientific). Ppard agonist GW501516 (1 μM; Sigma-Aldrich) was added to organoid cultures for the mice treated with the GW501516 diet. Organoids were imaged on days 1, 3, and 6 of culture unless otherwise specified.

### QUANTIFICATION AND STATISTICAL ANALYSIS

Statistical significance was determined by the unpaired Student *t*-test, Poisson test, or analysis of variance (one-way or two-way) with Bonferroni adjustments for all multiple comparisons. The significance of correlations was determined by the Spearman correlation coefficient. Survival time was calculated using the Kaplan-Meier method and compared between groups using the log-rank test. All tests were two-sided, and significance was defined as *P*<.05. Data were analyzed using SAS software v9.4 (SAS Institute) or GraphPad Prism v7.01 (GraphPad Software).

## References

1. Kamisawa T, Wood LD, Itoi T, et al. Pancreatic cancer. Lancet 2016;388:73–85.

2. Kanda M, Matthaei H, Wu J, et al. Presence of Somatic Mutations in Most Early-Stage Pancreatic Intraepithelial Neoplasia. Gastroenterology 2012;142:730–733.e9.

3. di Magliano MP, Logsdon CD. Roles for KRAS in Pancreatic Tumor Development and Progression. Gastroenterology 2013;144:1220–1229.

4. Nöthlings U, Wilkens LR, Murphy SP, et al. Meat and fat intake as risk factors for pancreatic cancer: the multiethnic cohort study. J Natl Cancer Inst 2005;97:1458–65.

5. Thiébaut AC, Jiao L, Silverman DT, et al. Dietary fatty acids and pancreatic cancer in the NIH-AARP diet and health study. J Natl Cancer Inst 2009;101:1001–11.

6. Gordon-Dseagu VLZ, Thompson FE, Subar AF, et al. A Cohort Study of Adolescent and Midlife Diet and Pancreatic Cancer Risk in the NIH-AARP Diet and Health Study. Am J Epidemiol 2017;186:305–317.

7. Hori M, Kitahashi T, Imai T, et al. Enhancement of carcinogenesis and fatty infiltration in the pancreas in N-nitrosobis(2-oxopropyl)amine-treated hamsters by high-fat diet. Pancreas 2011;40:1234–40.

8. Philip B, Roland CL, Daniluk J, et al. A high-fat diet activates oncogenic Kras and COX2 to induce development of pancreatic ductal adenocarcinoma in mice. Gastroenterology 2013;145:1449–58.

9. Khasawneh J, Schulz MD, Walch A, et al. Inflammation and mitochondrial fatty acid beta-oxidation link obesity to early tumor promotion. Proc Natl Acad Sci U S A 2009;106:3354–9.

10. Chang HH, Moro A, Takakura K, et al. Incidence of pancreatic cancer is dramatically increased by a high fat, high calorie diet in KrasG12D mice. PLoS One 2017;12:e0184455.

11. Dawson DW, Hertzer K, Moro A, et al. High-fat, high-calorie diet promotes early pancreatic neoplasia in the conditional KrasG12D mouse model. Cancer Prev Res (Phila) 2013;6:1064–73.

12. Luo Y, Yang Y, Liu M, et al. Oncogenic KRAS Reduces Expression of FGF21 in Acinar Cells to Promote Pancreatic Tumorigenesis in Mice on a High-Fat Diet. Gastroenterology 2019;157:1413–1428 e11.

13. Nadella S, Burks J, Al-Sabban A, et al. Dietary fat stimulates pancreatic cancer growth and promotes fibrosis of the tumor microenvironment through the cholecystokinin receptor. Am J Physiol Gastrointest Liver Physiol 2018;315:G699–G712.

14. Takahashi M, Hori M, Ishigamori R, et al. Fatty pancreas: A possible risk factor for pancreatic cancer in animals and humans. Cancer Sci 2018;109:3013–3023.

15. Liu Y, Colby JK, Zuo X, et al. The Role of PPAR-delta in Metabolism, Inflammation, and Cancer: Many Characters of a Critical Transcription Factor. Int J Mol Sci 2018;19.

16. Beyaz S, Mana MD, Roper J, et al. High-fat diet enhances stemness and tumorigenicity of intestinal progenitors. Nature 2016;531:53–8.

17. He TC, Chan TA, Vogelstein B, et al. PPARdelta is an APC-regulated target of nonsteroidal antiinflammatory drugs. Cell 1999;99:335–45.

18. Gupta RA, Tan J, Krause WF, et al. Prostacyclin-mediated activation of peroxisome proliferator-activated receptor delta in colorectal cancer. Proc Natl Acad Sci U S A 2000;97:13275–80.

19. Abdollahi A, Schwager C, Kleeff J, et al. Transcriptional network governing the angiogenic switch in human pancreatic cancer. Proc Natl Acad Sci U S A 2007;104:12890–5.

20. Pedchenko TV, Gonzalez AL, Wang D, et al. Peroxisome Proliferator–Activated Receptor β/δ Expression and Activation in Lung Cancer. American Journal of Respiratory Cell and Molecular Biology 2008;39:689–696.

21. Muller-Brusselbach S, Komhoff M, Rieck M, et al. Deregulation of tumor angiogenesis and blockade of tumor growth in PPARbeta-deficient mice. EMBO Journal 2007;26:3686–98.

22. Peters JM, Gonzalez FJ, Müller R. Establishing the Role of PPARβ/δ in Carcinogenesis. Trends Endocrinol Metab 2015;26:595–607.

23. Zuo X, Xu W, Xu M, et al. Metastasis regulation by PPARD expression in cancer cells. JCI Insight 2017;2:e91419.

24. Coleman JD, Thompson JT, Smith RW, et al. Role of Peroxisome Proliferator-Activated Receptor and B-Cell Lymphoma-6 in Regulation of Genes Involved in Metastasis and Migration in Pancreatic Cancer Cells. PPAR Research 2013;2013:11.

25. Choi YJ, Roberts BK, Wang X, et al. Effects of the PPAR-delta agonist MBX-8025 on atherogenic dyslipidemia. Atherosclerosis 2012;220:470–6.

26. Risérus U, Sprecher D, Johnson T, et al. Activation of Peroxisome Proliferator–Activated Receptor (PPAR)δ Promotes Reversal of Multiple Metabolic Abnormalities, Reduces Oxidative Stress, and Increases Fatty Acid Oxidation in Moderately Obese Men. Diabetes 2008;57:332–339.

27. Fan W, Waizenegger W, Lin CS, et al. PPARδ Promotes Running Endurance by Preserving Glucose. Cell Metabolism 2017;25:1186–1193.e4.

28. Hingorani SR, Petricoin EF, Maitra A, et al. Preinvasive and invasive ductal pancreatic cancer and its early detection in the mouse. Cancer Cell 2003;4:437–50.

29. Grizzi F, Fiorino S, Qehajaj D, et al. Computer-aided assessment of the extra-cellular matrix during pancreatic carcinogenesis: a pilot study. J Transl Med 2019;17:61.

30. Georgoudaki AM, Prokopec KE, Boura VF, et al. Reprogramming Tumor-Associated Macrophages by Antibody Targeting Inhibits Cancer Progression and Metastasis. Cell Rep 2016;15:2000–11.

31. Movahedi K, Laoui D, Gysemans C, et al. Different tumor microenvironments contain functionally distinct subsets of macrophages derived from Ly6C(high) monocytes. Cancer Res 2010;70:5728–39.

32. Zuo X, Deguchi Y, Xu W, et al. PPARD and Interferon Gamma Promote Transformation of Gastric Progenitor Cells and Tumorigenesis in Mice. Gastroenterology 2019;157:163–178.

33. Ying H, Dey P, Yao W, et al. Genetics and biology of pancreatic ductal adenocarcinoma. Genes Dev 2016;30:355–85.

34. Hou P, Kapoor A, Zhang Q, et al. Tumor Microenvironment Remodeling Enables Bypass of Oncogenic KRAS Dependency in Pancreatic Cancer. Cancer Discov 2020;10:1058–1077.

35. Li BH, Garstka MA, Li ZF. Chemokines and their receptors promoting the recruitment of myeloid-derived suppressor cells into the tumor. Mol Immunol 2020;117:201–215.

36. Ying H, Kimmelman AC, Lyssiotis CA, et al. Oncogenic Kras maintains pancreatic tumors through regulation of anabolic glucose metabolism. Cell 2012;149:656–70.

37. Shao J, Sheng H, DuBois RN. Peroxisome proliferator-activated receptors modulate K-Ras-mediated transformation of intestinal epithelial cells. Cancer Res 2002;62:3282–8.

38. Ugel S, De Sanctis F, Mandruzzato S, et al. Tumor-induced myeloid deviation: when myeloid-derived suppressor cells meet tumor-associated macrophages. The Journal of Clinical Investigation 2015;125:3365–3376.

39. Marvel D, Gabrilovich DI. Myeloid-derived suppressor cells in the tumor microenvironment: expect the unexpected. The Journal of Clinical Investigation 2015;125:3356–3364.

40. Veglia F, Perego M, Gabrilovich D. Myeloid-derived suppressor cells coming of age. Nature Immunology 2018;19:108–119.

41. Gabrilovich DI, Ostrand-Rosenberg S, Bronte V. Coordinated regulation of myeloid cells by tumours. Nat Rev Immunol 2012;12:253–68.

42. Wynn TA. Myeloid-cell differentiation redefined in cancer. Nat Immunol 2013;14:197–9.

43. Mantovani A, Marchesi F, Malesci A, et al. Tumour-associated macrophages as treatment targets in oncology. Nat Rev Clin Oncol 2017;14:399–416.

44. Stromnes IM, Brockenbrough JS, Izeradjene K, et al. Targeted depletion of an MDSC subset unmasks pancreatic ductal adenocarcinoma to adaptive immunity. Gut 2014;63:1769–81.

45. Kaur K, Chang H-H, Topchyan P, et al. Deficiencies in Natural Killer Cell Numbers, Expansion, and Function at the Pre-Neoplastic Stage of Pancreatic Cancer by KRAS Mutation in the Pancreas of Obese Mice. Frontiers in Immunology 2018;9.

46. Bayne LJ, Beatty GL, Jhala N, et al. Tumor-derived granulocyte-macrophage colony-stimulating factor regulates myeloid inflammation and T cell immunity in pancreatic cancer. Cancer Cell 2012;21:822–35.

47. Fleming V, Hu X, Weber R, et al. Targeting Myeloid-Derived Suppressor Cells to Bypass Tumor-Induced Immunosuppression. Front Immunol 2018;9:398.

48. Ugel S, De Sanctis F, Mandruzzato S, et al. Tumor-induced myeloid deviation: when myeloid-derived suppressor cells meet tumor-associated macrophages. J Clin Invest 2015;125:3365–76.

49. Huang H, Liu Y, Daniluk J, et al. Activation of nuclear factor-kappaB in acinar cells increases the severity of pancreatitis in mice. Gastroenterology 2013;144:202–10.

50. Daniluk J, Liu Y, Deng D, et al. An NF-kappaB pathway-mediated positive feedback loop amplifies Ras activity to pathological levels in mice. J Clin Invest 2012;122:1519–28.

51. Wei D, Wang L, Yan Y, et al. KLF4 Is Essential for Induction of Cellular Identity Change and Acinar-to-Ductal Reprogramming during Early Pancreatic Carcinogenesis. Cancer Cell 2016;29:324–338.

52. Barak Y, Liao D, He W, et al. Effects of peroxisome proliferator-activated receptor delta on placentation, adiposity, and colorectal cancer. Proc Natl Acad Sci U S A 2002;99:303–8.

53. Liu Y, Deguchi Y, Tian R, et al. Pleiotropic Effects of PPARD Accelerate Colorectal Tumorigenesis, Progression, and Invasion. Cancer Res 2019;79:954–969.

54. Charafe-Jauffret E, Tarpin C, Bardou VJ, et al. Immunophenotypic analysis of inflammatory breast cancers: identification of an ‘inflammatory signature’. J Pathol 2004;202:265–73.

55. Yan Z, Gao J, Lv X, et al. Quantitative Evaluation and Selection of Reference Genes for Quantitative RT-PCR in Mouse Acute Pancreatitis. Biomed Res Int 2016;2016:8367063.

