## Supplemental Figures 1-5 with legends for "Ppard Is Essential in Acceleration of Pancreatic Ductal Adenocarcinoma Development by High-Fat Diet in Mutant *Kras* Mice"

Figure S1

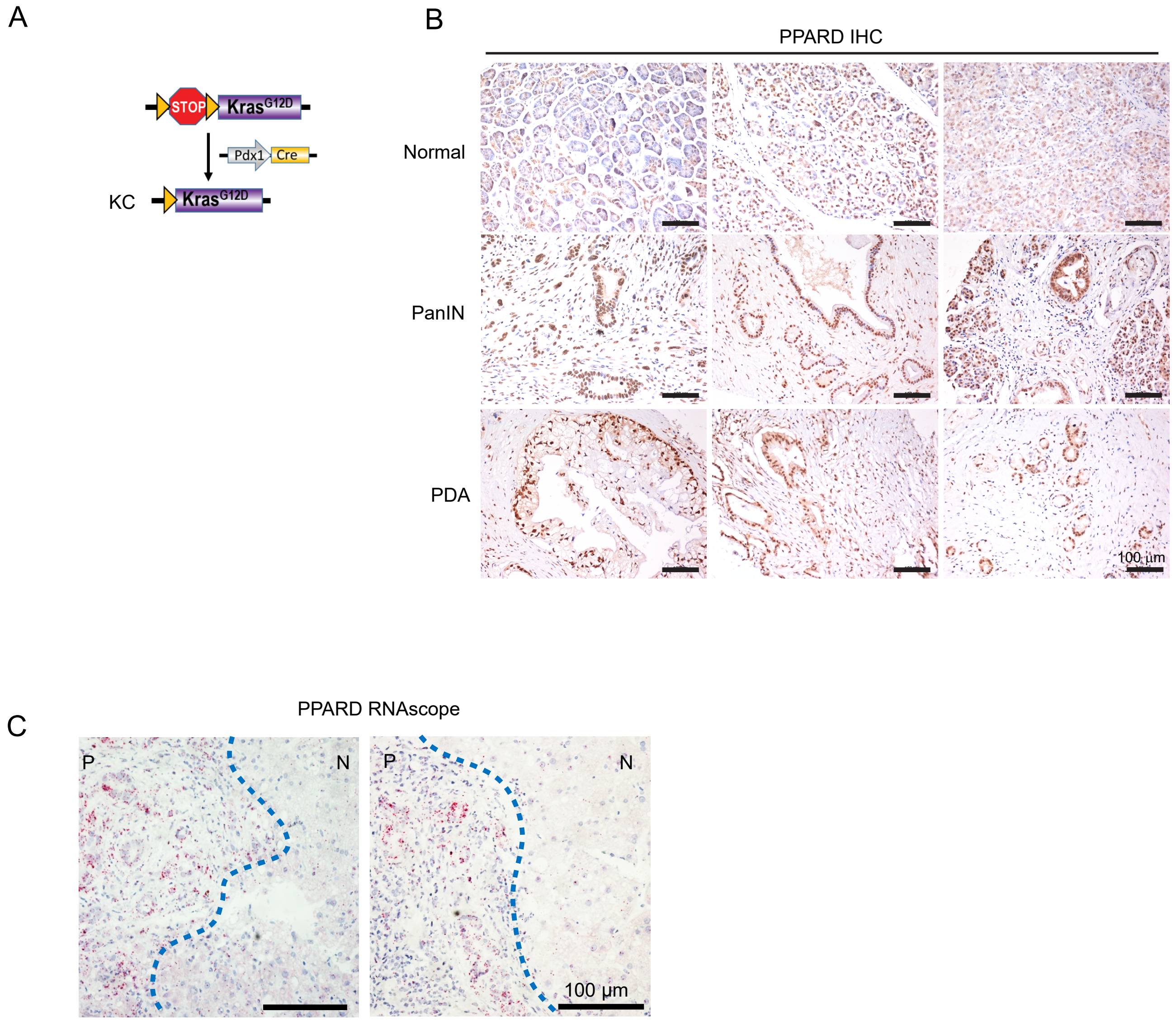


**Figure S1. PPARD is upregulated in pancreatic cancer. Related to Figure 1.**
**(A)** Schematic diagram for the generation of KC mice.
**(B)** Representative images of PPARD immunochemistry staining for human pancreatic tissue microarray samples with normal, PanIN, and PDAC areas.
**(C)** Representative images of PPARD RNAscope in situ hybridizations for human biopsied pancreatic tissue samples, including PanINs (P) and pancreatic normal areas (N).

Figure S2

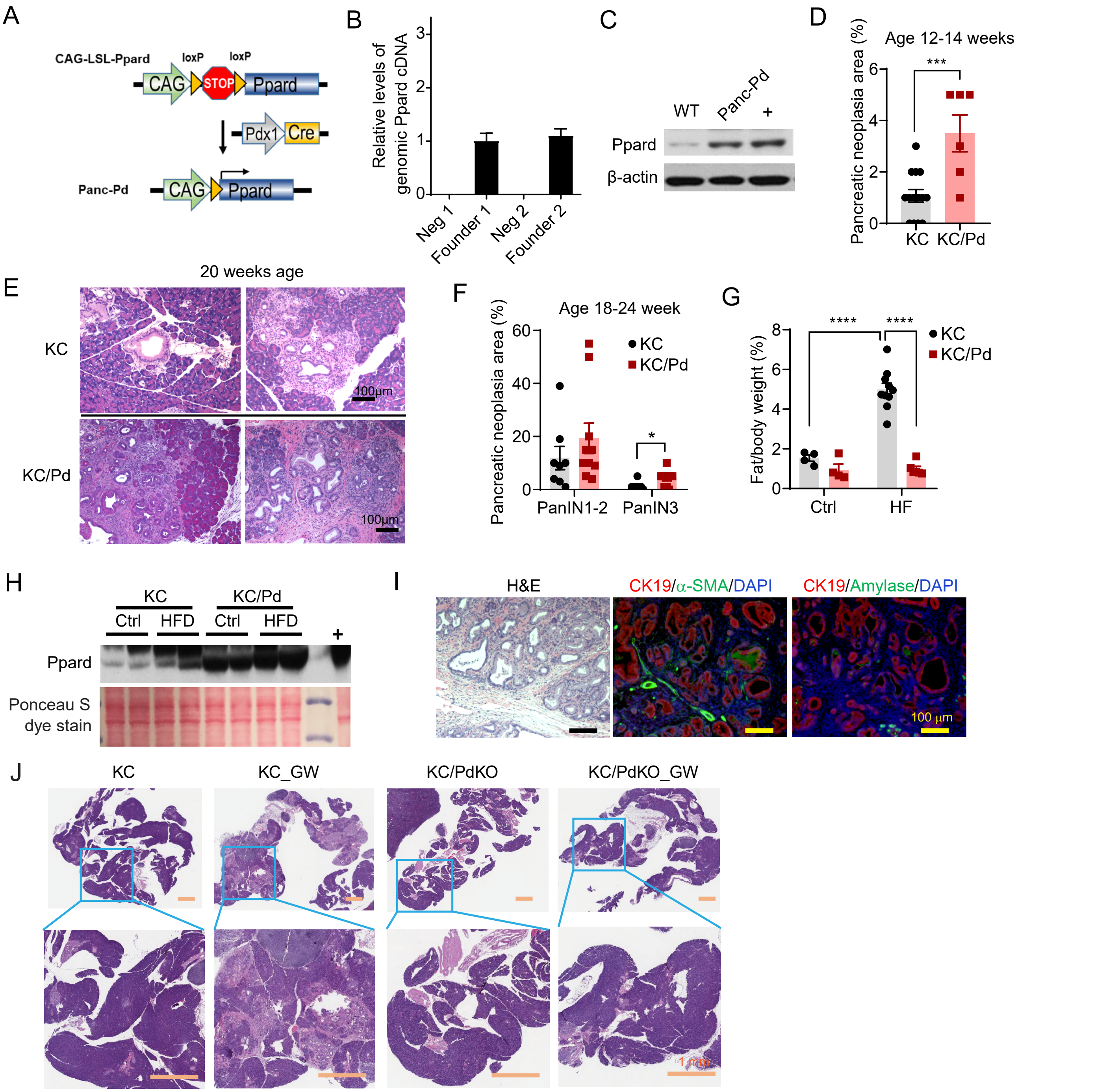


**Figure S2. Hyperactivation of Ppard promotes pancreatic tumorigenesis in KC mice. Related to Figures 2 and 3.**
**(A)** Schematic diagram of the generation of Panc-Pd mice. CAG-LSL-Ppard transgenic mice were bred with *Pdx1-Cre* mice to generate *Pdx1-Cre*, CAG-LSL-*Ppard* mice, designated as Panc-Pd mice.
**(B, C)** Characterization of Panc-Pd mice. The relative levels of genomic Ppard cDNA **(B)** in tails, measured by qPCR, and Ppard protein levels in mouse pancreata **(C)**, measured by Western blot in Panc-Pd mice. "+" indicates positive control of Ppard.
**(D)** Comparisons of percentages of pancreatic neoplasia areas between KC and KC/Pd mice at age 12-14 weeks (n=6-12 mice per group).
**(E, F)** Histologic characterization of the pancreata in KC and KC/Pd mice at age 20 weeks. **(E)** Representative images of H&E staining for the indicated mice are shown. **(F)** Comparisons of the percentages of pancreatic neoplastic areas shown as PanIN 1-2 and PanIN 3 per mouse for the indicated mice (n=8-9 mice per group).
**(G)** The weight ratio of abdominal fat to body per mouse for KC and KC/Pd mice fed a high-fat diet (HFD) or control (Ctrl) as described in **Fig. 2B-E**.
**(H)** Ppard protein expression in pancreata in KC and KC/Pd mice fed HFD) or Ctrl as described in **panel G**, measured by Western blot. Ponceau S dye–stained bands in membrane were evaluated as an internal reference. "+" indicates the Ppard+ control.
**(I)** Representative images of H&E staining and co-immunofluorescence staining of CK19 and α-SMA or CK19 and amylase in pancreata of KC/Pd mice fed a GW diet (50 mg/kg) for 20 days.
**(J)** Representative images of H&E staining in pancreata of KC and KC/PdKO mice fed the GW (50 mg/kg) or control (Ctrl) diet for 13 weeks as described in **Fig. 3H-L**.
For **D** and **F**, the data are mean ± SEM, *$P$ <.05, and ***$P$ <.001.

Figure S3

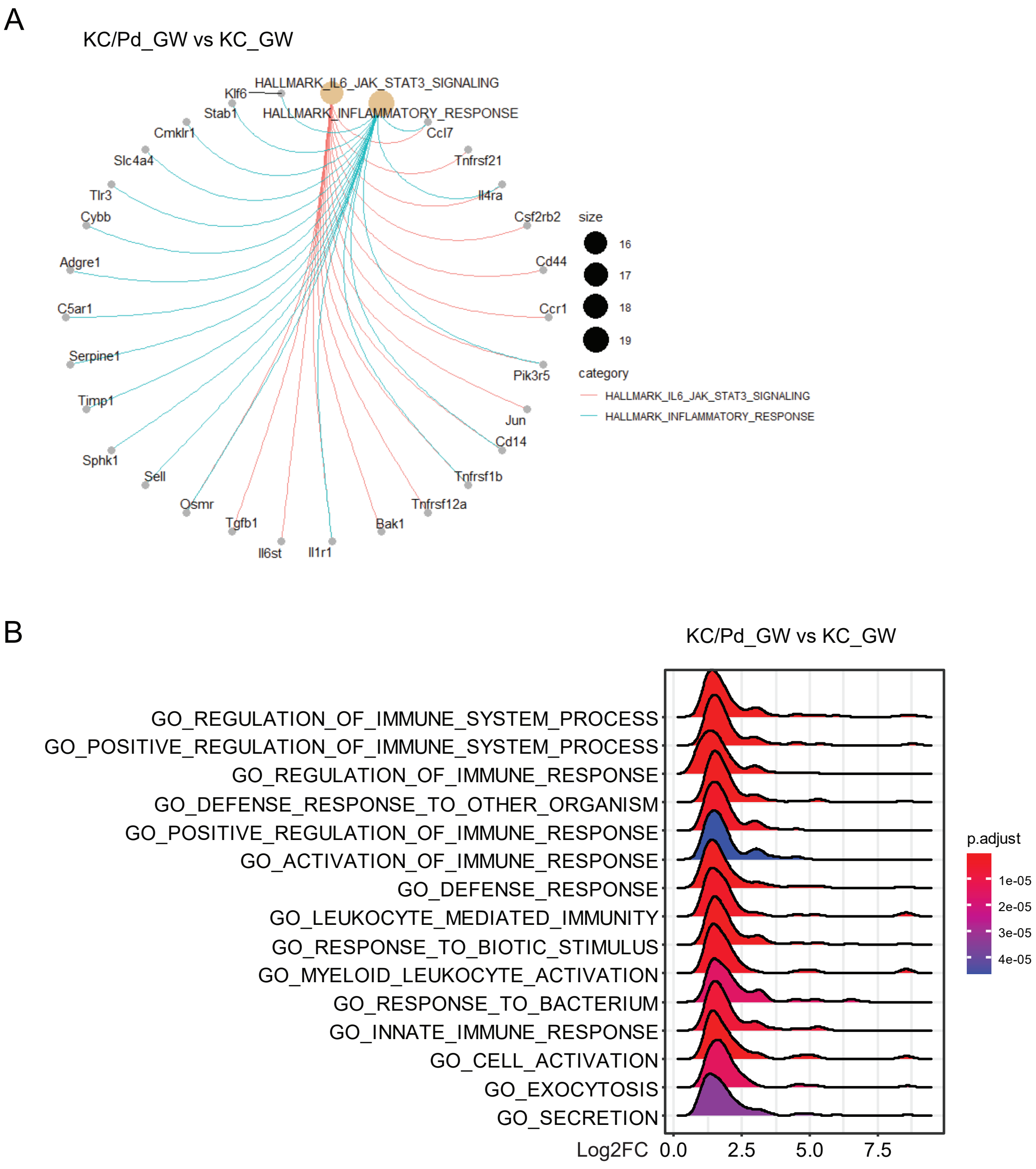


**Figure S3. Ppard hyperactivation drives inflammatory response–related pathways identified by GSEA. Related to Figure 4.**
**(A)** The Cnet plots of two top inflammatory response–related pathways using Hallmark gene sets, as described in **Fig. 4B**.
**(B)** GSEA results of KC and KC/Pd mice fed a GW diet for 3 days (KC_GW versus KC/Pd_GW), as described in **Fig. 4** using R package clusterProfiler, pAdj Cutoff=0.05, gene sets = MSigDB category "GO gene sets". The ridgeline plots of top 15 enriched pathways.

Figure S4

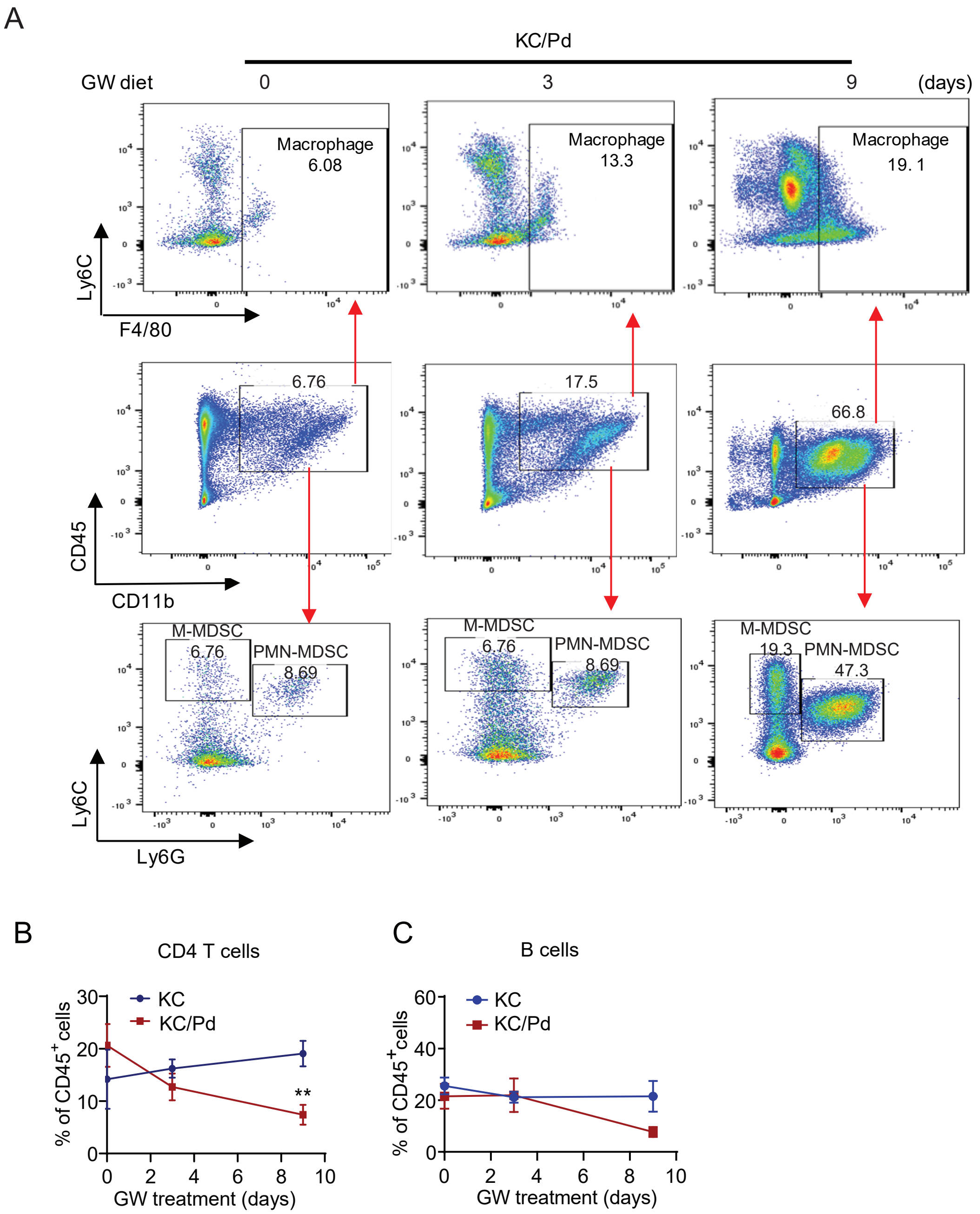


**Figure S4. Ppard hyperactivation orchestrates an immunosuppressive microenvironment to promote pancreatic tumorigenesis in KC mice. Related to Figure 5.**

**(A-C)** KC and KC/Pd mice at 6-8 weeks were fed the GW (50 mg/kg) for 0, 3, or 9 days (n=3-5 per group), and then euthanized. Their pancreata were examined for infiltrating immune cells assessed by flow cytometry as described in **Fig. 5A-H**. **(A)** Representative flow cytometry gating image results for pancreas-infiltrating macrophages ($CD45^+CD11b^+F4/80^+$), M-MDSCs ($CD45^+CD11b^+Ly6C^{hi}Ly6G^-$) and PMN-MDSCs ($CD45^+CD11b^+Ly6C^+Ly6G^+$). **(B, C)** Percentage of $CD3^+CD4^+$ T cells **(B)** and B cells ($B220^+$) **(C)** among pancreas-infiltrating CD45+ cells for the indicated mouse groups. Data are mean ± SEM; ***P* <.01.

### Figure S5

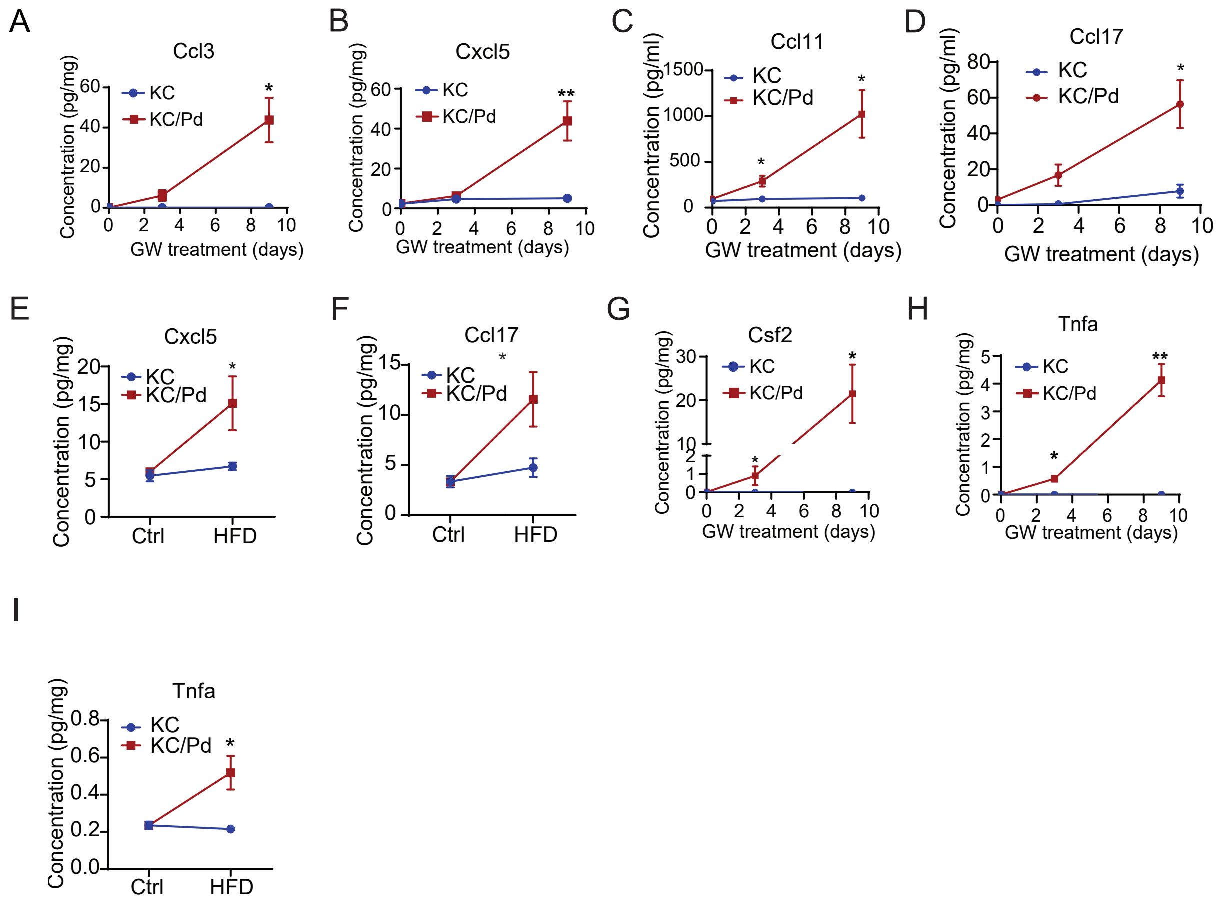


**Figure S5. PPARD hyperactivation increases various chemokines and cytokines to establish an immunosuppressive microenvironment. Related to Figure 6.**
**(A-I)** KC and KC/Pd mice at 6-8 weeks were fed the GW diet (50 mg/kg) for 0, 3, or 9 days, or a high-fat diet (HFD) or control (Ctrl) diet for 13 weeks, and then euthanized. The pancreata protein lysates were examined for a panel of chemokines and cytokines by Biolegend's LEGENDplex bead-based immunoassays as described in Fig. 6A-L (n=3-5 per group). **(A-F)** Chemokine protein concentration of pancreata from KC or KC/Pd mice fed the GW diet for Cxcl5 **(A)**, Cxcl5 **(B)**, Ccl11 **(C)**, and Ccl17 **(D)**, or fed either the HFD or Ctrl diet for Cxcl5 **(E)**, and Ccl17 **(F)** are shown. Cytokine protein concentrations of pancreata from KC or KC/Pd mice fed the GW diet for Csf2 **(G)** and Tnfa **(H)** or fed either the HFD or Ctrl diet for Tnfa **(I)** are shown. Data are mean ± SEM; **P* <.05, ***P* <.01.
